# Multiscale spatial analysis implicates chromosomal metaloops in gene patterning across the *Drosophila* brain

**DOI:** 10.64898/2026.09.02.748911

**Authors:** Aleena L. Patel, Achuthan Raja Venkatesh, Tohn Borjigin, Xiao Li, Michael S. Levine, Alistair N. Boettiger

## Abstract

Scores of chromosome-scale loops, or metaloops, arise in the *Drosophila* brain, but their spatial organization and relationship to neural gene expression patterns remain unclear. Here, we used multiplexed Optical Reconstruction of Chromatin Architecture (ORCA) to examine the multiscale spatial organization of metaloops in cross-sections of 100s of larval and adult *Drosophila* brains. We find metaloops form preferentially in the central regions of the brain, where they nucleate the formation of metadomains, characterized by the intermingling of distal topologically associating domains (TADs). At the sub-cellular scale, metaloops tend to arise towards the nuclear center, and multiple metaloops in the same cell have a preference to form hubs (3 or more contacts). Each brain nucleus generally harbors only a few loops or hubs. An in-depth analysis of the hub centered on *DIP-epsilon*, a synaptic wiring gene, identified a three-way metadomain that brings together the *DIP-epsilon* TAD; a distal TAD carrying a paralog of *DIP-epsilon*, *DIP-zeta*; and a putative regulatory TAD, across 3 Mb. This metadomain adopts distinct conformations depending on gene expression; cells expressing *DIP-epsilon* or *DIP-zeta* show preferential interactions between the TAD carrying the corresponding gene and the putative regulatory TAD. We posit that the neuron-specific formation of different subsets of metadomains might coordinate the expression of diverse combinations of synaptic wiring genes underlying complex brain architecture.

## Introduction

Chromatin loops bypass the constraints of the linear genome in order to co-localize discontiguous genomic segments within a nucleus. Linking enhancers and target gene promoters via chromatin loops is a strategy used during embryogenesis to deploy gene transcription in precise spatiotemporal patterns^1–6^. Variations in 3D genome folding further regulate gene expression choices during neuronal cell maturation^7–10^. For instance, tuning cohesin-driven extrusion of chromosomal loops determines which protocadherin (Pcdh) gene promoter contacts a distal enhancer, a mechanism that allows for diversity in Pcdh expression among mammalian neurons^11,12^. In olfactory sensory neurons, multi-chromosomal hubs colocalize olfactory receptor (OR) genes and their cognate enhancers to mediate singular OR gene choice^13,14^. In the *Drosophila* central nervous system, many megabase-scale chromosomal contacts emerge at genes implicated in axon guidance and synapse organization distributed throughout the genome^15–17^. Given the complexity of gene expression patterns involved in neural circuit specification and assembly, genome organization may vary significantly from cell to cell in nervous systems. Specialization of genome organization in any nervous system has been under explored, due in part to challenges handling the diversity of cell types and of preserving the spatial arrangement of neurons, which is critical to the function of a nervous system.

Across metazoans, gene-regulatory enhancer-promoter (E-P) contacts are typically found in sub-megabase scale topologically associating domains (TADs)^18^. However, recent studies in *Drosophila* brains, using Micro-C^19^, identified a surprising set of contacts in bulk data linking genes and prospective enhancers across distances up to the length of entire chromosome arms ^15^. These ultra-long-range loops were called “metaloops”, and were substantially enriched for promoters of genes with major roles in neural cell maturation. It remains unclear if these metaloops are a pan-neuronal level of genome organization, or a superposition of loops that exist in distinct cell types. The majority of metaloops connect regions of the genome occupied by the transcription factors CTCF and/or Cp190, which are broadly expressed in the brain^15,20,21^. Many metaloops are weakened upon perturbation to these factors – which could suggest a pan-neuronal character^16,18^. The genes that are hypothesized to be linked to distant enhancers by metaloops encode cell adhesion molecules, neurotransmitter receptors, ion channels, and other factors specializing neuronal function and known to be expressed in subsets of cells^22^. Non-uniform metaloop associations across the brain may be important to limit gene expression to the right neural cell types. An unbiased survey of *in situ* metaloop organization would reveal how differentiated neural 3D genomes are among single cells.

Metaloop interactions detected by Micro-C co-occur with enriched contacts among sequences surrounding the anchor, a feature called “metadomains”, rather than appearing only as point-to-point interactions^15^. Visualizing how gene-regulatory sequences, normally confined to TADs, associate with their long range partners in metadomains could shed light on mechanistic models of metadomain assembly and regulation of genes housed in them.

To address the limitations of existing bulk technologies, and to map cell-specific variation in heterogeneous nervous system tissue, we imaged folded chromatin segments *in situ* ^4,23^. Here, we applied Optical Reconstruction of Chromatin Architecture (ORCA)^24^ to cryosectioned larval and adult brains. We first mapped spatially variable contacts of metaloop anchors genome-wide, across the developing larval brain, and discovered non-uniform patterns of metaloop assembly. We detected multi-metaloop clusters, and found that clustered sequences are spatially arranged within the nuclear volume. We then traced entire TADs and flanks surrounding three distal metaloop anchors, which we found to form hubs in this first analysis. We found that two TADs housing metaloop-proximal genes compete for contact with a third distal TAD in transcriptionally active cells, in which these genes guide synaptic connections. Our multiscale imaging supports a view in which metadomains contribute to neural function by co-localizing linearly isolated genomic loci, allowing for higher-order 3D interactions to differentially regulate neural genes that ultimately shape functional circuits underlying complex behaviors^25^.

## Results

### The multiscale spatial organization of metaloops *in situ*

The *Drosophila* larval brain has been an exemplary model in which to study neural circuitry because of its sophisticated structural complexity set up by a diversity of well-mapped cell types^26,27^. We performed an extensive imaging study to probe the heterogeneity and functional role of megabase-scale genome organization (**Fig. 1a**) within the nuclei of these diverse cells distributed across cryosectioned brain tissue. Towards this end, we located the positions of sequences that co-localize to form metaloops using unique sequence tags targeted to 73 independent genomic regions (metaloop anchors, **Supplementary Table 1, Supplementary Fig. 1a**), which we detected with fluorescent oligos using ORCA^24^ within ∼100 individual third instar larval brains imaged in the same experiment. Several metaloop anchors are proximal to the transcription start site (TSS) of a gene, and therefore are named promoter (P) anchors, while others have been classified as intergenic (I)^15^. We targeted anchors involved in all 58 metaloops detected by Micro-C and found 49 could be robustly detected with ORCA, as well as 10 short-range anchor interactions within TADs (called intraTAD loops) (**Supplementary Fig. 1b**). We named the loops according to the convention introduced in Mohana, Dorier & Li et al^15^.

**Fig. 1.**
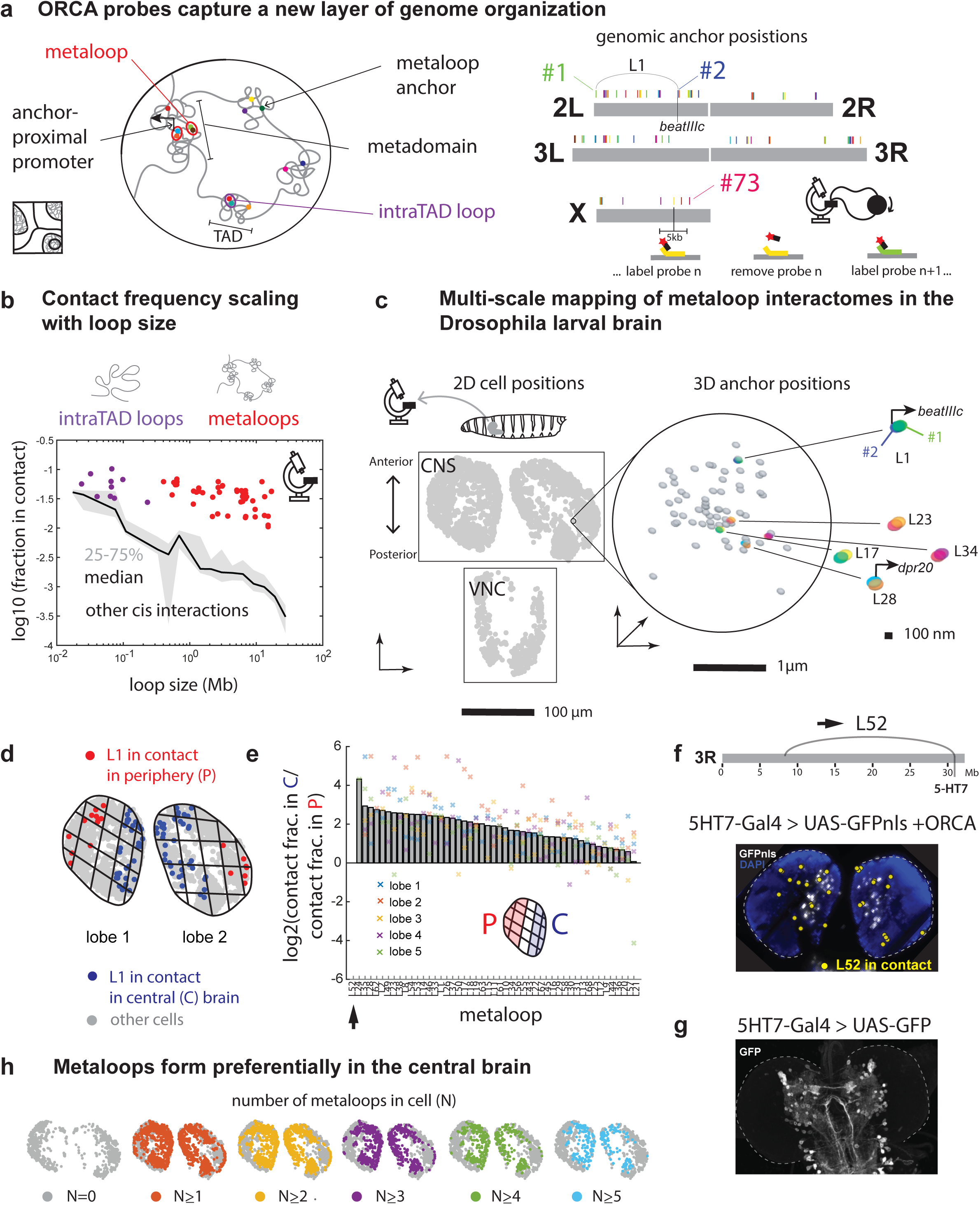
Multiscale organization of metaloops *in situ*. **a,** 73 DNA FISH probes targeted to metaloop anchors distributed across the *Drosophila* genome capture a layer of genome organization that connects TADs into metadomains. Several of these probes are proximal to important neural genes, e.g. *beatIIIc*. **b,** Contact frequency, measured as the fraction of cells with anchors less than 150 nm apart, for metaloops (red), intraTAD loops (purple), and all other loops connected by probes on the same chromosome (black line showing median and shaded region showing 20-75% confidence interval), of various sizes. **c,** The 2D positions of cells in brain tissue cross sections and 3D positions of ORCA probes within individual nuclei are captured at once. An example cross-section of a 3rd instar larval brain and zoom-in to a single nucleus are shown. The anchors of metaloops that are “in contact” are colored in the inset, and ORCA probes proximal to neural gene promoters are labeled (*beatIIIc* and *dpr20*). **d,** An example cross section of a larval CNS with the cells with the metaloop L1 in contact in the peripheral (P) pixels highlighted red, and in the central (C) pixels blue. The other cells are in grey. **e,** The log2 fold change in frequency of each metaloop in P versus C, in 5 example cross sections, ranked by the mean. L52 is the top ranking metaloop (arrow). **f,** L52 (arrow) connects *5-HT7* to a very distal metaloop anchor on chromosome 3R. The image shows GFPnls driven by a Gal4 driver labeling 5-HT7 expression in an example cross section of the larval CNS, counterstained with DAPI. The locations of cells with L52 formed are shown. **g,** The 5-HT7 Gal4 driving GFP labeling neuronal circuitry in the CNS. **h,** 2D cell positions in an example CNS cross section colored by the number (N) of metaloops in each cell (N = 0, more than 1, 2, 3, 4, or 5; grey, red, yellow, purple, green, blue respectively).

In our images, we defined metaloops “in contact” based on the euclidean distances between ORCA probes. After accounting for potential imaging artefacts (see Methods) we retained 79,732 cells, with a high probe detection efficiency (**Supplementary Fig. 2)**. In these cells, we observed over 400 million inter-metaloop anchor 3D distances. The distances between anchor pairs that constitute metaloops were distributed bimodally, suggesting that not every metaloop is “in contact” in every cell (**Supplementary Fig. 3a)**. We calculated contact probabilities based on a 3D distance threshold and produced a heatmap that correlated well with Micro-C data binned to the size of ORCA probes (Pearson’s *R* = 0.867) (**Supplementary Fig. 3b,c**). While the probability of contact between anchors in pairs that are not predicted to form a metaloop, or form intraTAD loops, fell with increasing genomic separation, metaloop contacts were frequently detected with minimal effect of genomic separation (**Fig. 1b**). We conclude that distance-based assessment of “contacts” captures the enrichment of specific interactions leading to metaloops and intraTAD loops in cell types of the nervous system, originally detected by Micro-C, while enabling a new investigation of metaloop interactomes in single cells, spatially resolved.

Since populations of cells with metaloop anchors in contact and not in contact both exist, metaloops could occur stochastically throughout the tissue, or within distinct cell populations that harbor subsets of metaloops. Our multi-scale and spatial microscopy data (**Fig. 1c**) permitted deeper investigation of metaloop organization in order to address this question.

### Metaloops preferentially form in the central brain

To analyze how various metaloops differed in their organization across the brain, we selected cross-sections that captured the optic lobes (OL) and central brain (CB), which comprise the larval central nervous system (CNS), and aligned each cross section so that it could be segmented into discrete pixels along the anterior (An) - posterior (Post) and peripheral (P) - central (C) axes (**Fig. 1d, Supplementary Fig. 4**). All 48 metaloops were more likely to occur in the central pixels than in the peripheral ones. This bias was substantially greater for some loops, e.g. L52 was ∼16-fold C/P enriched, while L21 showed a <2-fold difference (**Fig. 1e**). The spatial biases of metaloops along the An-Post axis were more modest (<2-fold) (**Supplementary Fig. 5**). Interestingly, the highly spatially biased metaloop (L52) has one anchor proximal to a key serotonin receptor, 5-HT7, needed only in mature cells ready to form synapses^28–30^. Using a Gal4 driver to label 5-HT7 positive nuclei, we mapped these nuclei and found them also enriched in the central region of the brain (where L52 preferentially formed) (**Fig. 1f**), localization that is consistent with prior tracing of serotonin-sensitive cells^31^ (**Fig. 1g**).

To analyze how brain regions differed in their propensity to form metaloops, we mapped the total number of metaloops formed per cell (**Fig. 1h**). Most cells harbored less than 5 simultaneously formed metaloops (**Supplementary Fig. 6a**). We found that cells lacking metaloops were enriched in the periphery, while those with more metaloops were increasingly concentrated toward the central brain. The spatial gradients of cells as a function of number of metaloops (N=1, 2, 3, 4, and 5) were steeper along the P-C axis than along the An-Post axis (**Supplementary Fig. 6b-d**), mirroring what we observed on a loop-by-loop basis. We searched further for patterns of metaloop co-occurrence by applying dimensionality reduction to the single-cell contact data (**Supplementary Fig. 7a**). This produced 28 distinct clusters, with the major groups representing cells with no loops at all, various single loops, or more than 3 loops (**Supplementary Fig. 7b,c**). While there was no systematic separation within the multi-loop cluster, the multi-loop cells appeared more abundant in the central regions of the CNS cross-sections than the no-loop or single-loop cells (**Supplementary Fig. 7d**).

The pattern of metaloop abundance along the peripheral-central brain axis, and correlation of individual loops with the expression of mature neuronal factors follows a known gradient of neural differentiation along the P-C axis^26,27,32^. We posit that the higher order genome folding mechanisms that lead to abundant metaloops are influenced or driven by neural maturation, which in many cases may coincide with spatially patterned activation of anchor-proximal neural genes, like 5-HT7. Within multi-loop cells, metaloops potentially directly affect one another, for example by shortening chromosomes, further shaping the diverse metaloop interactomes distributed among mature neural cell-types.

### Multiple metaloop anchors frequently coalesce in hubs due to loop stacking

We next focused on cells in the multi-loop cluster to investigate metaloop interactions. We considered where metaloop anchors are situated in the genome relative to one another, as this positioning could influence whether one metaloop sterically hinders or facilitates a neighboring metaloop on the same chromosome. For instance, two metaloops that are separated on opposite ends of the same chromosome might form independently of one another. When loops are nested, on the other hand, the formation of one loop could facilitate the formation of the other by folding up the chromosome and bridging much of the intervening genomic distance (**Fig. 2a**). When two loops are crossed, they may antagonize one another if the formation of one loop pulls the anchor of another nearby metaloop away from its partner. Two loops may share a common anchor, a configuration previously called stacked loops^33^. In principle the underlying interactions could form competitively, leading to single loops, or cooperatively, leading to metaloop anchor hubs. While a roughly equal proportion of metaloop pairs that are separated, nested, or crossed were found to be either correlated or anticorrelated with one another, all instances of stacked metaloop pairs were correlated (**Fig. 2b**), reflecting a tendency of metaloop anchors to form hubs (**Fig. 2c, Supplementary Fig. 8a,b**), rather than independent or competing pairwise contacts.

**Fig. 2.**
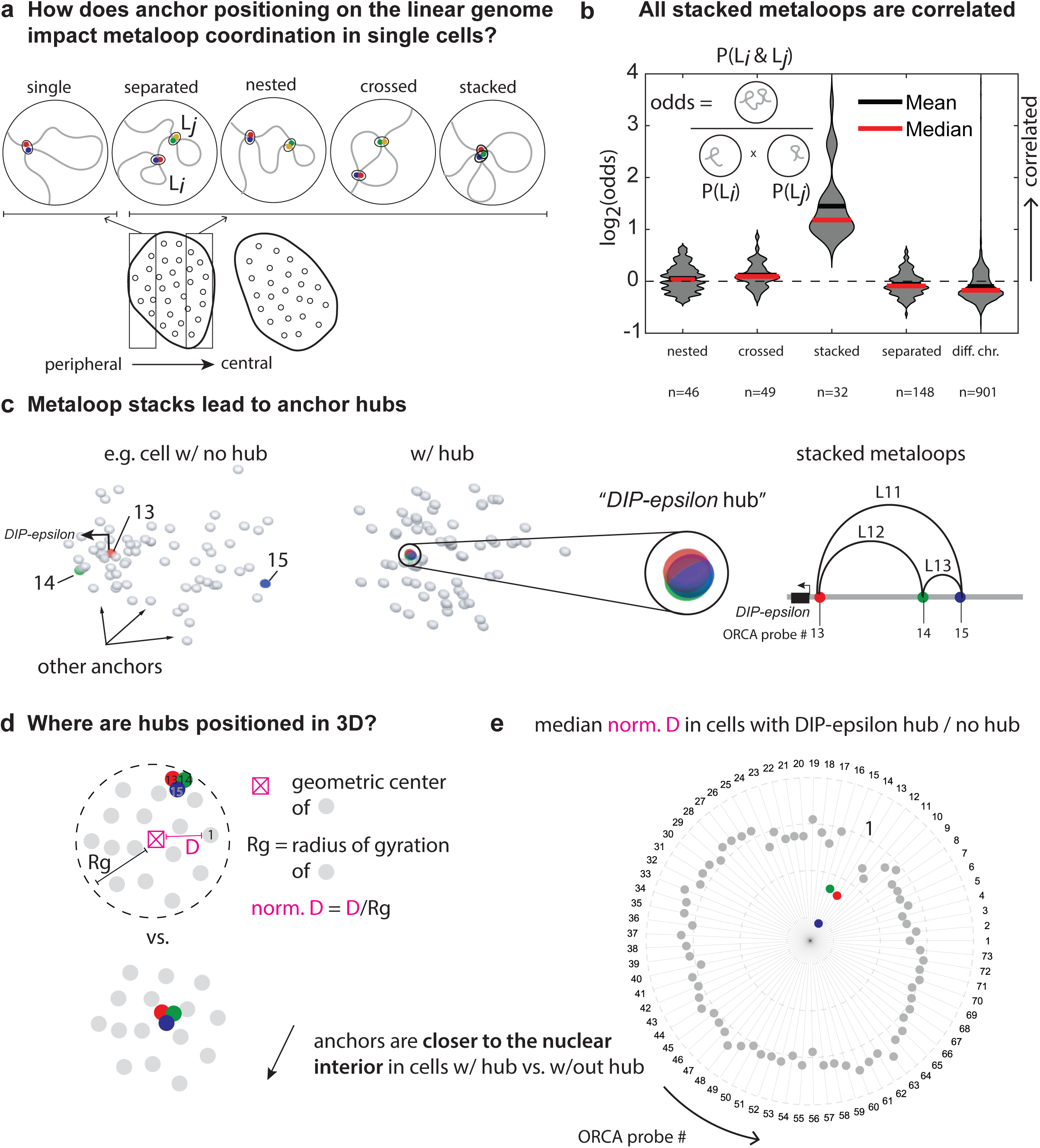
Stacked metaloops coalesce into multi-way hubs towards the nuclear interior. **a,** Pairs of metaloops, e.g. L*_i_* and L*_j_*, can be separated, nested, crossed, or stacked depending on where their anchors lie on the genome. Single loops are generally more prevalent in the periphery. **b,** The log_2_ odds that pairs of metaloops are both in contact at the same time, for pairs classified as nested, crossed, stacked, or having no overlap because they are separated on the same chromosome, or on different chromosomes. Values greater than 0 (dotted line) indicate metaloops that are correlated with one another. The number of pairs (n) belonging to each category are shown. **c,** Example plots of the 3D positions of ORCA probes labeling anchors in a hub (left) and not in a hub (right). The hub includes the anchor proximal to the gene *DIP-epsilon*. This *DIP-epsilon* hub occurs when 3 metaloops L11, L12, and L13 stack together. **d,** Hub positioning is assayed by measuring the distance “D” between ORCA probes and the geometric center of anchors (excluding those labeled by ORCA probes 13,14, and 15). “D” is normalized by the radius of gyration of ORCA probe positions filling the volume of each nucleus. **e,** A circle plot showing the ratio of the median normalized distances to the geometric center (norm. D) for cells with the *DIP-epsilon* hub versus cells with anchors 13, 14 and 15 dispersed (no hub). ORCA probe numbers corresponding to anchors are labeled on the periphery of the plot, and values at 1 fall on the dotted circle are labeled.

### Metaloops and metaloop hubs preferentially arise in the nuclear interior

We observed that when metaloop anchors were associated in hubs, they were often nearer the center of the nucleus rather than at the periphery (example hub shown in **Fig. 2c**). To quantify this centering effect, we computed the distance of hub-associated anchors from the centroid of all 73 anchors. We compared this distance to that of all other anchors not participating in the hub (**Fig. 2d**). This analysis revealed a significant displacement of hub anchors towards the center (**Fig. 2e**, **Supplementary Fig. 8c-e**). In cell clusters with only one metaloop, the anchors in contact were also closer to the nuclear interior, suggesting that centering is not unique to hubs, but a feature of metaloop formation in general (**Supplementary Fig. 8f**).

To look at how other anchors responded to hub formation, we considered differences in contact and distance maps built from cells with and without hubs (**Supplementary Fig. 9**). Contact frequencies between anchors not participating in the hub were largely unchanged, indicating that the formation of a hub is not accompanied by many other formed metaloops (**Supplementary Fig. 9**). However, when we considered differences in 3D distances between anchors, we found the median distances between anchors participating in the hub and all other anchors to be decreased without necessarily leading to a contact (**Supplementary Fig. 9**). This reflects the tendency of anchors in hubs to be more centrally located in the nucleus, and hence also closer to any other part of the genome.

The preference that specific sets of metaloops stack and form hubs away from the nuclear periphery does not support a model in which solely shortening of chromosomes facilitates higher-order organization of metaloops. Rather, transcriptionally active chromatin at metaloop anchors may be stabilized at regions in the central nucleus, such as a nuclear speckle^34,35^ or TF clusters^36,37^, or expelled from repressive compartments at the periphery^38–40^.

When distal anchors form hubs, it is possible that the flanking chromatin around these anchors also changes structure, for example stretching out to form new contacts with the flanks of hub partners, or compacting away to be isolated from them. To better understand the behavior of the flanking chromatin, we turned to a targeted approach, focusing on on a hub identified by our pan-anchor labeling approach that involved an anchor proximal to the promoter of *DIP-epsilon*, a downstream anchor near *DIP-zeta* ∼2.5 Mb away, and a third anchor ∼3 Mb away (**Fig. 2c**). Both *dpr*-interacting protein (DIP) genes play important roles in neural wiring^41–43^.

### Hub formation alters spatial organization of distal TADs as they intermesh

We extended our probes around each anchor participating in the *DIP-epsilon* hub, to nearby promoters (like *DIP-zeta*), candidate enhancers (ATAC sensitive regions) and TAD borders, to test alternate models of how these regions interact (**Fig. 3a**, **Supplementary Fig. 10a**). We tiled the *DIP-epsilon* and *DIP-zeta* gene-containing TADs and flanking DNA (chr2L 6.3-6.45Mb and chr2L 8.95-9.15Mb respectively), as well as the TAD and flanking DNA containing the third anchor (chr2L 9.4 to 9.55Mb), using 92 uniquely barcoded probes (**Fig. 3b**). The probes spanned 5 anchors leading to 4 metaloops (**Fig. 3b, Supplementary Fig. 10a**). Tracing data from larval brains reproduced the pairwise metaloop contacts detected in our larval pan-metaloop analysis (**Supplementary Fig. 10b**). Contact maps generated from ORCA traces in adult brains showed well-delineated TADs and quantitatively stronger metaloop interactions, consistent with prior Micro-C, and we focused our analysis in this tissue (**Fig. 3b, Supplementary Fig. 10c,d**). We refer to average contacts between distal domains (typically represented as heatmaps) as metadomains (**Fig 3b**).

**Fig. 3.**
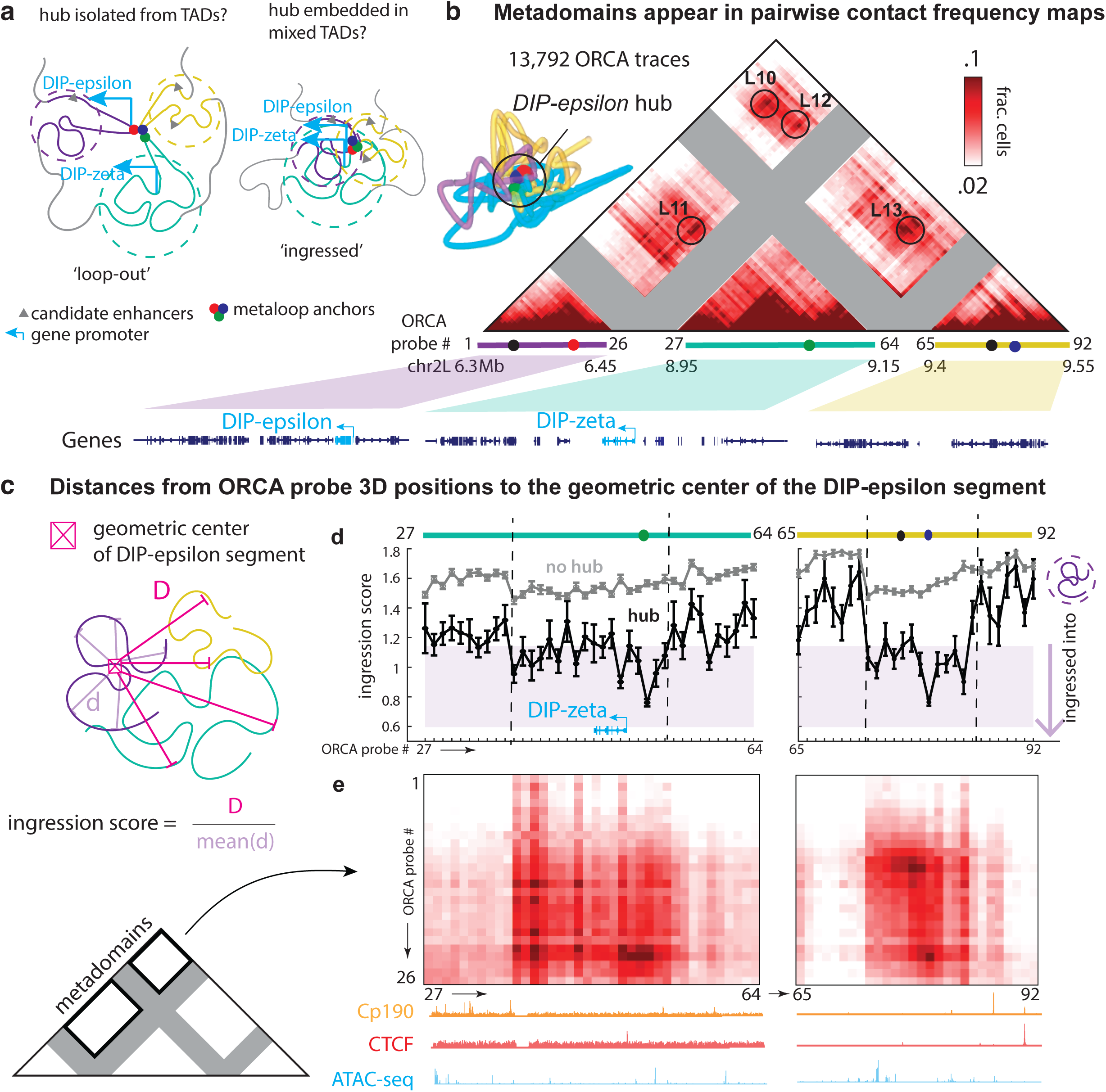
Hub formation alters spatial organization of distal TADs as they intermesh. **a,** Metaloop anchors could make contact outside of the volume occupied by its local surrounding chromatin (‘loop-out’ model), or while embedded, leading to ingressed TADs (‘ingressed’ model). Regulatory elements distal to anchors in a hub would be excluded from contacting one another in the loop-out model, and more likely to make contact in an ingressed model. **b,** An example trace of flanking chromatin around the *DIP-epsilon* hub, and a contact frequency map built from 13,792 such traces. The pairwise interactions corresponding to the metaloops L10, L11, L12, and L13 are circled. ORCA probes traced chr2L 6.3-6.45 Mb, which spans the *DIP-epsilon* gene, chr2L: 8.95-9.15 Mb, which spans the *DIP-zeta* gene, and a region downstream, chr2L: 9.4-9.55 Mb. Gene tracks are shown below the contact map. **c,** Distances to the geometric center of the *DIP-epsilon* containing segment were normalized by the mean of all distances between probes in the *DIP-epsilon* segment and their geometric center and reported as an ingression score. **d,** The mean and s.e.m. of the ingression scores for cells with (grey) and without the *DIP-epsilon* hub (black). The shaded region marks the 25-75% confidence interval for all pooled distances between *DIP-epsilon* region probes and their geometric center. **e,** Contact maps showing averaged interactions between the *DIP-epsilon* segment and the two segments downstream of it. ChIP-seq tracks for Cp190 and CTCF (data from ^44^) and ATAC-seq tracks (data from ^61^) for these distal segments are shown.

Analysis of these traces revealed distinct spatial organization of TADs depending on whether or not the anchors were engaged in a hub. When in a hub, the *DIP-epsilon*-proximal anchors were positioned closer to the geometric center of the *DIP-epsilon* region compared to traces without the hub, inconsistent with the loop-out hypothesis (**Fig. 3a**, **Supplementary Fig. 11a**). This repositioning of anchors in a hub toward the interior of its own region was apparent when measuring probe distances to the geometric centers of the *DIP-zeta* and 3rd anchor-TAD regions as well (**Supplementary Fig. 11b,c**).

Hub-containing cells also exhibited a complex pattern of inter-TAD organization. Of all elements in the TAD, the *DIP-zeta* anchor was closest to the center of the *DIP-epsilon* region (most ingressed, lower ingression score) (**Fig. 3c,d**). However, several other elements throughout the *DIP-zeta* TAD were as close on average to the center of the *DIP-epsilon* region as the average *DIP-epsilon* element was to its own center, also inconsistent with the loop-out hypothesis (**Fig. 3d, left**). Similar behavior was observed when considering ingression of the 3rd anchor-TAD into the *DIP-epsilon* region (**Fig. 3d, right**) and the other combinations of interacting TADs (**Supplementary Fig. 11d,e**). Moving out from the hub anchors, we observed, not a steady increase in ingression, but a fluctuating change followed by sharp jumps at TAD boundaries marked by binding of the insulators CTCF or Cp190^44^ (**Fig. 3d,e**). Similarly fluctuating ingression patterns, as observed for the *DIP-epsilon* region, occurred in our measurements of the ingression into the *DIP-zeta* and 3rd anchor-TAD regions upon hub formation (**Supplementary Fig. 11d,e**). The sharp changes in ingression across the TAD borders suggest that boundaries delineate not only insulated local cis interactions, but also reduced long-range (trans) interactions (**Supplementary Fig. 11f**).

In our ingression analyses, we calibrated the statistical degree of ingression by comparing the ingression scores of elements distal to the region whose geometric center was mapped, to the ingression scores of elements within that region. For example, the shaded region in **Fig. 3d** indicates the 25-75% confidence interval for all pooled ingression scores between *DIP-epsilon* region probes and their geometric center. Scores for the ingression of elements in the *DIP-zeta* and 3rd anchor-TAD regions that fall within the shaded box (**Fig. 3d**) indicate substantive intermingling of these distal chromatin segments with the *DIP-epsilon* region, defining interactions spatially, rather than from averaged contacts (metadomains) (**Fig. 3e**).

We expected that the elements of each TAD would expand to accommodate ingression of the distal TADs, but instead we found they compact (**Supplementary Fig. 11g**). Given that corresponding TADs ingress when hubs occurred, rather than the anchors looping out of the TADs to form hubs, this tightening of TAD structures when they join the hub was surprising. This observation is, however, consistent with a view in which the intra-TAD contacts and inter-TAD contacts are both facilitated by a shared set of bridging molecules, so having more of the latter also increases contact in the former.

### *DIP-epsilon* and *DIP-zeta* expressing cells form distinctly organized metadomains

To further investigate how intermingling of TADs in the fly brain relates to the differential regulation of neural wiring genes, we performed additional ORCA experiments in adult brains in which either *DIP-epsilon* or *DIP-zeta* positive cells were marked by GFP using a GAL4 driver. We elected a *DIPepsilon*/*zeta*-GAL4>UAS-GFPnls approach to reliably distinguish the gene-expressing from silent nuclei (see **Fig. 4a,b, Supplementary Fig. 12)**. We obtained 6,720 *DIP-epsilon* positive traces to compare to 11,005 *DIP-epsilon* negative traces, and 2,600 *DIP-zeta* positive traces to compare to 23,006 *DIP-zeta* negative traces. We compared contact maps generated from each sub-population of traces obtained in the same experiment.

**Fig. 4.**
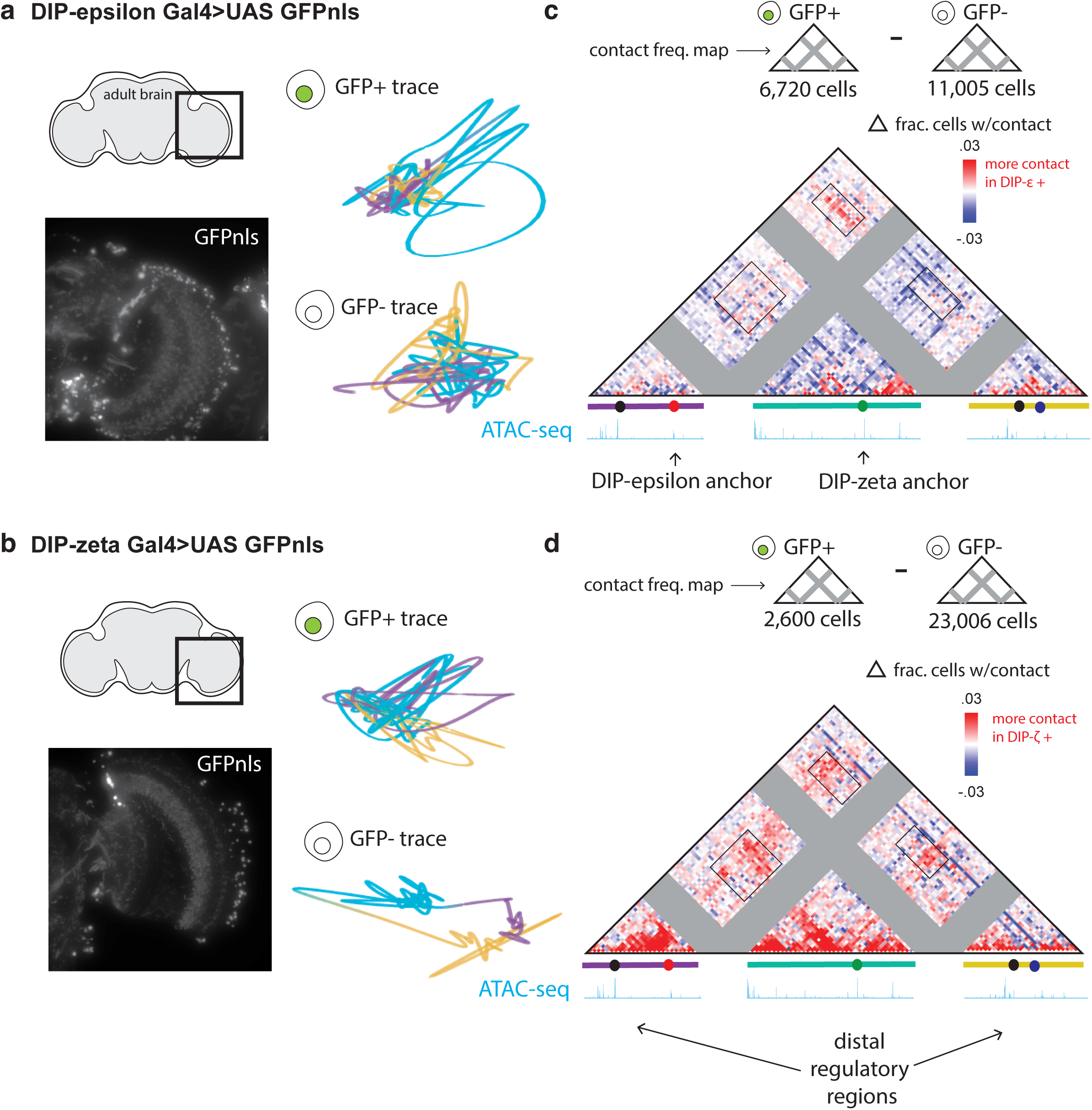
Pairwise contact differences in metadomain traces correlate with gene expression. **a,** An image of an adult *Drosophila* brain cryosection labeled with GFP marking *DIP-epsilon* expression using the Gal4/UAS system. Example traces of the 3 segments, also traced in Fig. 3, in a GFP+ and GFP-cell are shown. **b,** An image of an adult *Drosophila* brain cryosection labeled with GFP marking *DIP-zeta* expression using the Gal4/UAS system. Examples of GFP+ and GFP-traces are shown. **c,** Difference in pairwise contact frequency between *DIP-epsilon* expressing and silent cell populations. **d,** Difference in pairwise contact frequency between *DIP-zeta* expressing and silent cell populations. **c,d,** The boxes show metadomain boundaries. ATAC-seq peaks are shown, and in **d,** the arrows point out potential distal chromatin regions regulating *DIP-epsilon* and *DIP-zeta* expression.

Both *DIP-epsilon* and *DIP-zeta* expressing cells exhibited increased contact between the *DIP-epsilon* or *DIP-zeta* regions and the 3rd anchor-TAD compared to silent cells (**Fig 4c**,**d**). Because of this association of contact with expression, and the presence of candidate enhancer elements in the 3rd anchor-TAD, we hereafter refer to it as the “enhancer TAD”. In *DIP-epsilon* positive cells, the enhancer TAD is uniquely enriched in contact with the *DIP-epsilon* TAD, while showing little enrichment of contacts with the *DIP-zeta* TAD (**Fig. 4c**). Shorter average 3D distances between the probes tiling the *DIP-epsilon* and *DIP-zeta* TADs and the enhancer TAD also reflect these gene-dependent differences in contact (**Supplementary Fig. 13**).

In *DIP-zeta* expressing cells, additional interactions involving the *DIP-epsilon* TAD occurred. Pairwise interactions between a region upstream of the *DIP-epsilon* gene and the *DIP-zeta* gene were increased, suggesting that an additional regulatory element for *DIP-zeta* could sit in the *DIP-epsilon* region (**Fig. 4d, Supplementary Fig. 13e)**. Other pairs of paralogous genes have also been shown to be linked by long-range loops and shown to share an enhancer proximal to one of the genes^45,46^. For example, the cell adhesion gene, *sticks and stones (sns)* depends partly on distal enhancers in a TAD housing its paralogous gene, *hibris (hbs)*, linked by a metaloop^15,47,48^.

To further test the association of cross-TAD interactions on gene expression at additional loci, we examined the interactions between the TADs spanning *sns* and *hbs* in single cells of the larval brain. We labeled *sns*-expressing cells with a Gal4 driver^49^ and found *sns*-positive cells to be enriched for pairwise distal TAD contacts compared to *sns*-negative cells, similar to the increased co-localization of the *DIP-epsilon* and *DIP-zeta* TADs in *DIP-zeta* expressing cells (**Supplementary Fig. 14**). Since the *DIP-epsilon* and *DIP-zeta* TADs also contact regulatory chromatin in a third TAD distal to both genes, more complex metadomain configurations, such as competitive versus cooperative multi-TAD associations, could be additionally relevant for regulation of these genes.

### *DIP-epsilon* and *DIP-zeta* expression reflect competitive interactions among 3 distal TADs

*DIP-epsilon* and *DIP-zeta* play distinct roles in establishing connections among neurons, and are expressed in distinct patterns ^22,50–52^. *DIP-epsilon* and *DIP-zeta* expressing cells also both include multiple transcriptionally defined cell-types (**Supplementary Fig. 15**). To investigate whether selective long-range contacts could contribute to these differences, we considered the possible configurations of interactions among the three TADs in single cells (**Fig. 5a**). We considered two TADs to be “interacting” if at least 1 cross-TAD contact exists (**Fig. 5b**). We then compared the probabilities of finding cells in each state among active cells versus silent cells measured in the same experiment (**Fig. 5c**). The configuration in which all three TADs interact was most abundant in expressing and silent populations, consistent with our prior observations of anchor hub formation. The expression-dependent fold changes in TAD-TAD interaction state revealed that this configuration was mildly enriched in both *DIP-epsilon* and *DIP-zeta* expressing cells (**Fig. 5d**).

**Fig. 5.**
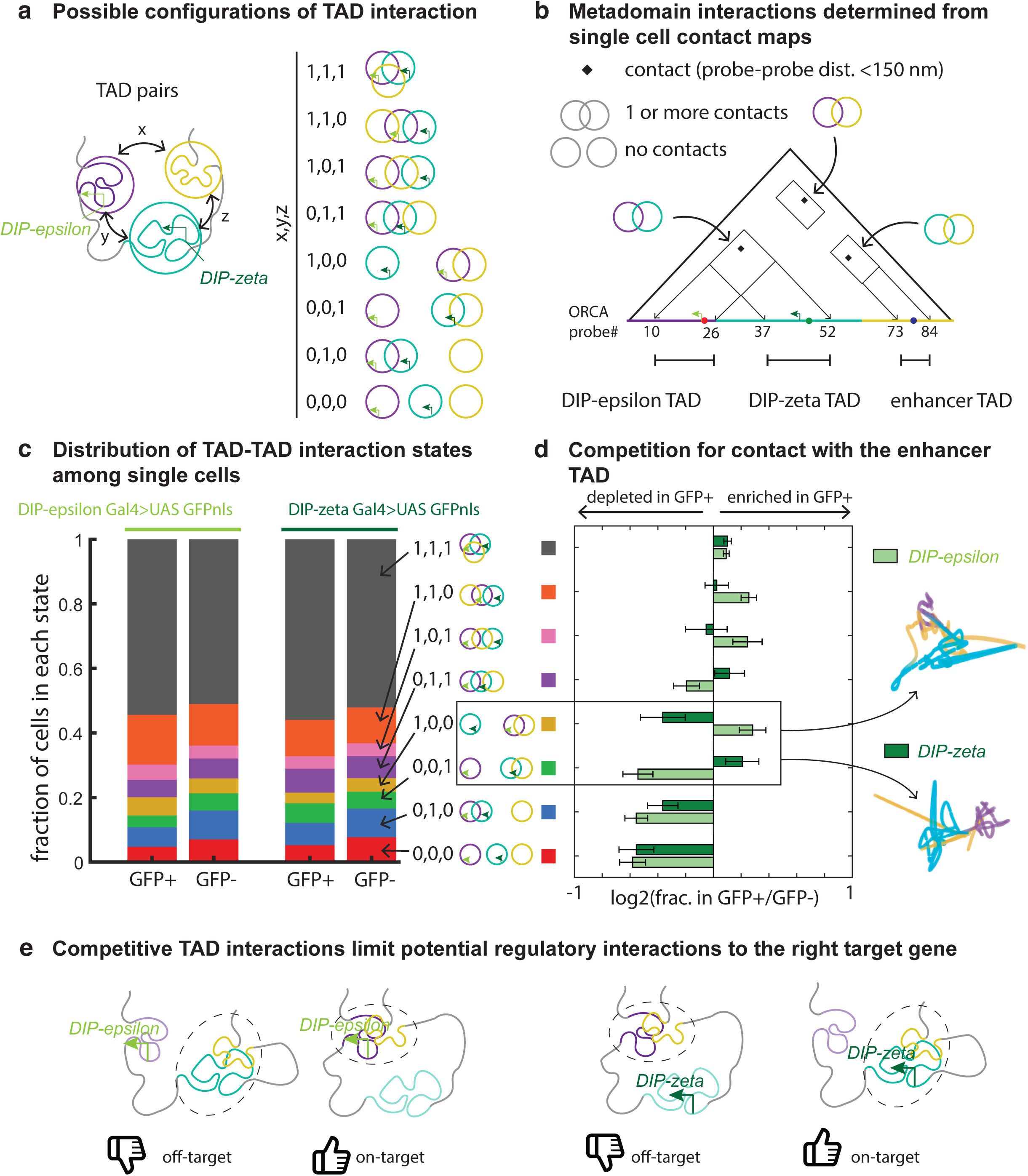
Competitive interactions underlie distinct gene expression states in the adult *Drosophila* brain. **a,** TAD pairs (x-x,x-y, and y-z) can interact in 8 possible configurations. **b,** TAD-TAD interactions states were measured in single cells. An interaction was counted if at least one contact was found between the probes tiling two TAD regions (indicated by the ORCA probe ranges). The TAD distal to both the *DIP-epsilon* and *DIP-zeta* containing TADs is named the “enhancer” TAD **c,** The proportions of cells with each configuration in the *DIP-epsilon* expressing and silent, and *DIP-zeta* expressing and silent populations. **d,** The log2 fold-change in abundance of each state in GFP+ versus GFP-cells, where GFP marks either *DIP-epsilon* or *DIP-zeta* expressing cells. Example traces of the highly competitive configurations that are enriched in the *DIP-epsilon* or *DIP-zeta* expressing cells are shown. **e,** A schematic showing the competitive TAD interactions in 3-way metadomains. When *DIP-epsilon* (light green) is expressed, its TAD is more likely exclusively interacting with the enhancer TAD. When *DIP-zeta* (dark green) is expressed, its TAD is more likely exclusively interacting with the enhancer TAD. This competition model limits which promoter accesses regulatory chromatin in a distal enhancer region.

However, among the cells in which any two of the TADs interact while the third is isolated, contacts between the *DIP-zeta* TAD and the enhancer TAD were favored in *DIP-zeta* expressing cells while disfavored in *DIP-epsilon* expressing cells, suggesting competitive regulation. Similarly, contacts between the *DIP-epsilon* TAD and the enhancer TAD were favored in *DIP-epsilon* expressing cells while disfavored in *DIP-zeta* expressing cells (**Fig. 5d**). We also noticed additional more nuanced competitive interactions, leading to the reduction of *DIP-epsilon* contacts with the enhancer TAD in *DIP-zeta* expressing cells, within contact maps generated from cells harboring 3-way TAD interactions (**Supplementary Fig. 16**). These analyses show that, in subsets of cells, competitive switches in metadomain contacts limit potential long-range regulatory interactions to the right target gene (**Fig. 5e**). These switch-like interactions within 3D genome structures, demonstrated by our single-cell metadomain tracing, may contribute to the spatial expression patterns of the cell adhesion genes in the brain, and thus play a key role in establishing correctly assembled circuits.

## Discussion

Here, we have produced spatial maps of long-range, gene-regulatory 3D genome interactions in single cells distributed throughout the *Drosophila* brain. We performed targeted labeling of sequences connected in metaloops, a recently discovered pattern of genome folding that links many neural gene promoters with distal, potential regulatory elements^15^. We found that metaloops formed non-uniformly across the CNS of the maturing larval brain, in a similar peripheral to central spatial pattern exhibited in the rate of differentiation of neurons, which are most mature at this stage in the central brain^26,27,32^. We hypothesize this spectrum of structural states reflects an increased reliance on long-range metaloops during neural differentiation.

Mammalian genomes also shift from relying on more proximal regulatory contacts to more distal ones throughout neural development. For example, high resolution maps from embryonic stem cells, neural precursor cells, and cortical neurons increase in loop size and in the strength of long range compartment interactions^53^. These include some examples of increased dependence on more long-range enhancers, such as at the gene *Sox2*^53,54^, or distinctive formation of multi-megabase scale, and even trans contacts, among OR genes and their enhancer clusters^14^. Single-cell analyses of genome structure using Dip-C and Pop-C in the human and mouse brain also uncovered strengthening of long-range contacts from early to late development^55^. We hypothesize that improved sensitivity to single-cell methods applied to animals with complex brains will continue to reveal sub-populations of cells with additional layers of genome folding enabling cis-regulatory elements to act at much longer-range, on a wider set of gene targets.

By correlating 3D genome structure with gene expression in adult *Drosophila* brains, we found that diverse metadomain configurations involving both cooperative and competitive interactions among TADs are distributed differently in transcriptionally active versus inactive cells expressing metaloop-proximal genes. These cell-type specific contact patterns in differentiated brain tissue contrast the frequently cell-type invariant patterns found earlier in embryogenesis. While the early *Drosophila* embryo exhibits numerous, specific long-range loops, including loops between paralogous genes (e.g. *scyl* and *chrb* or *kni* and *knrl)*, and other distal regulatory elements and associated genes (e.g. *Scr* and its blastoderm enhancer) ^45,46,56^, such loops have largely proved to be common among cells transcribing and not transcribing the associated genes^8,57,58^. Faster differentiation in embryos, which occurs in hours, compared to the days it takes to build a complete nervous system, may explain this difference. Diversified folding of the genome in neurons may allow the genomic blueprint to be reshaped and read in the many ways needed to direct the diverse cell behaviors that build an organ as complex as the brain.

From traces of TADs housing the crucial neural wiring genes *DIP-epsilon* and *DIP-zeta*^50–52^, we surmise that such diverse configurations are important to reduce simultaneous contact between the *DIP-epsilon* and *DIP-zeta* promoters with common enhancers in a distal TAD and facilitate their distinct expression patterns throughout the mature brain^59^. For example, a recently studied sub-population of cells in the adult brain, which differentially express *DIP-epsilon* and *DIP-zeta,* are a pair of neurons chiefly responsible for the fly’s escape response from looming visual stimuli^50^. The competitive structural interactions in the genome that we have imaged may help ensure that neuronal cross-talk is avoided. Similar mechanisms may be used widely to set up the extraordinarily precise spatiotemporal transcriptional programs behind neural specification and circuit assembly in the nervous system^60^.

We propose a model (**Fig. 6**) of metadomain formation that starts with distal genome regions (anchors) meeting one another and making contact in a pairwise manner or in hubs, which preferentially happens towards the nuclear interior. We speculate that two anchors make contact first, leading to single metaloops, before additional anchors join, form hubs, and increase the number of metaloops present per cell, during neural differentiation. Surrounding the co-localized anchors, the flanking DNA becomes intermeshed up to TAD boundaries. We further speculate that additional cell-type-specific chromatin factors fine-tune the intermeshed segments in a cell-type-specific manner. Alternatively, it is possible that competition and feedback in the process of intermeshing results in distinct 3D structural configurations that ultimately contribute to distinct cell-types.

**Fig. 6.**
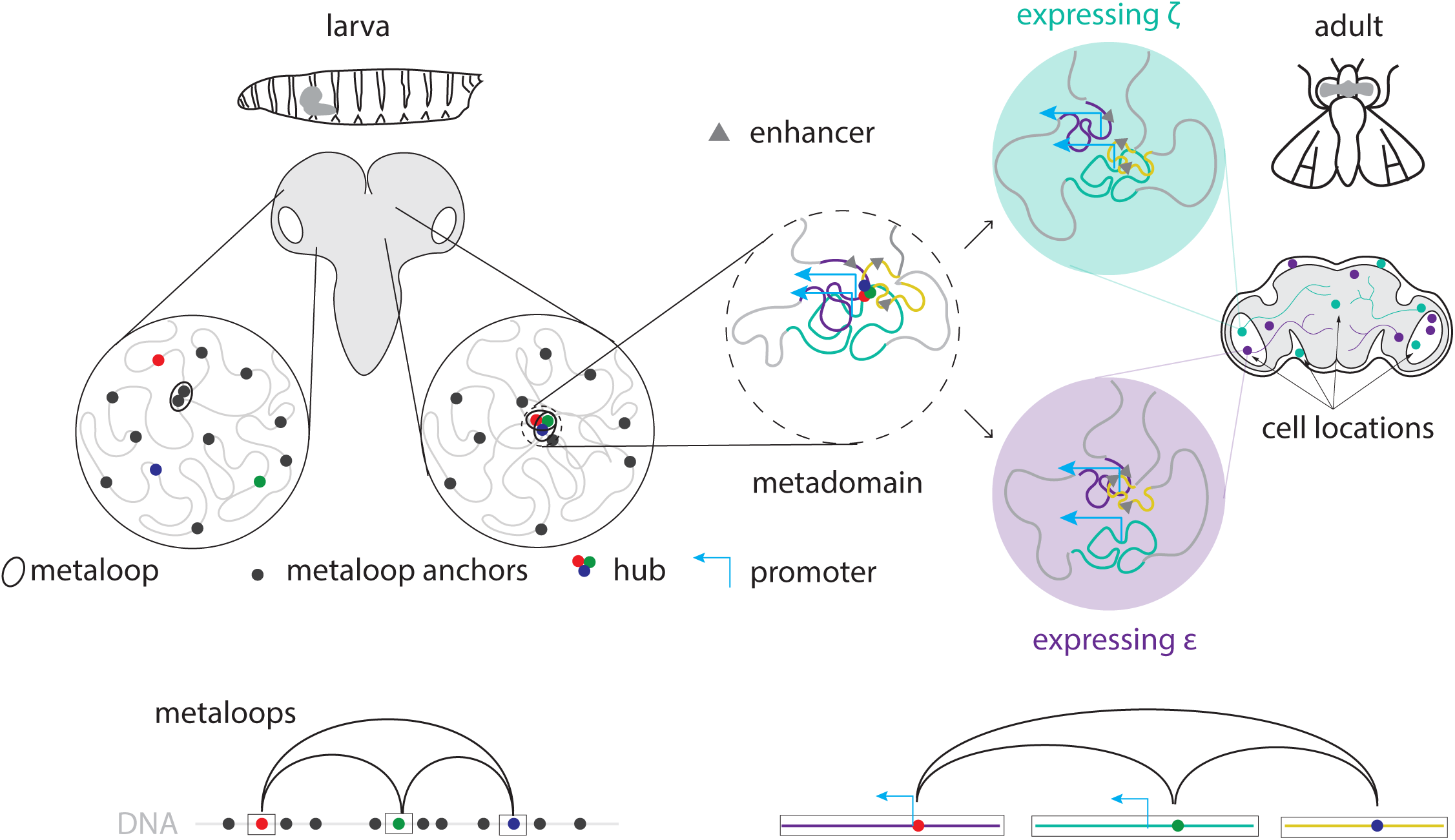
A summary model of metadomain assembly and function in the nervous system. A schematic summarizing the main findings of multiscale spatial imaging of chromosomal metaloops. Cells with no metaloops (dispersed anchors) or single metaloops (2 anchors in contact) were enriched in the peripheral regions of the developing larval brain, while cells with multiple metaloops forming hubs (3 or more anchors in contact) were concentrated towards the central brain. Targeted imaging of one hub showed 3 distal regions spanning two wiring genes (*DIP-epsilon* and *DIP-zeta*) to be enmeshed up to the boundaries of TADs. In the adult brain, cells expressing an anchor-proximal gene in the *DIP-epsilon* TAD (purple) or *DIP-zeta* TAD (teal) are scattered, and make synaptic connections with distinct partners during circuit assembly. Pairwise TAD-TAD interactions distinguish *DIP-epsilon* from *DIP-zeta* expressing cells in the adult brain.

Further investigation of the steps of metadomain assembly, and the potentially causative links between 3D genome structure, neuronal gene expression, and neuronal circuitry will require additional investigations in which spatiotemporally controlled genome folding perturbations are introduced. Although the incomplete list of candidate looping factors poses a challenge for designing such experiments, future investigations multiplexing ORCA with protein labeling and perturbation, in multiple stages of neurodevelopment, have strong promise to elucidate new mechanisms of neuronal gene control by 3D genome organization.

## Acknowledgements

The authors would like to thank members of the Boettiger and Nollmann labs for their critical feedback on the manuscript, as well as L. Luo (Stanford University) for gifting the *DIP-epsilon* Gal4 Drosophila line used in this study. This study was supported by a Helen Hay Whitney-AGBT Fellowship (A.L.P.), grants from the NIH, R01GM157292 (A.N.B) and R35GM118147 (M.S.L.), and a grant from the NSF, EF2022182 (A.N.B). A.L.P., A.N.B. and M.S.L. conceived the project. A.L.P. performed the experiments. A.L.P. and A.N.B. performed the data analysis. A.L.P. and A.N.B. wrote the manuscript with input from M.S.L., A.R.V., T.B., and X.L.

## Methods

### Sample collection and preparation for ORCA

#### Drosophila melanogaster transgenic lines

To label gene expression, we used Gal4 driver lines as follows. Bloomington Stock #46627 to label *5-HT7*, Bloomington Stock #90317 to label *DIP-zeta*, Bloomington Stock #76160 to label *sns*, and a DIP-epsilon-2A-Gal4 line gifted from the Liqun Luo lab at Stanford to label *DIP-epsilon*. These lines were crossed with a UAS_GFPnls line (Bloomington Stock #4775).

### Larval brain dissection & fixation

Larval brains were hand dissected from 3rd instar larvae harvested from the walls of culture vials. Larval bodies were inverted, exposing the larval brain while submerged in 1x PBS, and deposited in 4% PFA in 1x PBS. Larval brains attached to body walls were fixed overnight at 4C. After fixation, carcasses were washed 3x in 1x PBS to stop fixation, and transferred to a dissection dish. The larval brain was then isolated from the larval body wall, being careful to remove all imaginal discs and other non-brain tissue. Dissection was done in 1x PBS + .1% Tween-20 (PBST), and isolated larval brains were transferred to a tube containing the same buffer, which prevented sticking of the brain to the plastic tube. The PBST dissection buffer was then replaced with 100% methanol. Fixed brains stored in methanol were stored in -20C until preparation for cryosectioning.

### Adult brain dissection & fixation

Adult flies no older than 5 days were anesthetized on a carbon dioxide fly pad and decapitated. A random selection of male and female adults were included in each round of dissection. Adult heads were then transferred to a dissection dish containing PBST. The cuticle of the head was carefully removed to expose the adult brain using very sharp forceps. Tracheal tissue was also removed by hand. Individual adult brains were then manually transferred to 4% PFA in 1x PBST. As many brains as possible were dissected by hand within 45 mins, before the collection of dissected brains were left to fix at room temperature for 2 more hours, with gentle shaking. Some tubes were vortexed briefly to remove bubbles that caused brains to float to the top of the fixation solution. The fixation was stopped by washing 3x with PBST, and then PBST was replaced with methanol. Adult brains were stored in methanol at -20C until preparation for cryosectioning.

#### Cryosectioning

100% methanol was gradually replaced with PBST and brains were washed in PBST 3x. PBST was replaced with a fresh solution of 15% sucrose made in PBST, and stored at 4C until brains sank to the bottom of the tube. The 15% solution was then replaced with a 30% sucrose solution, and brains were stored in this solution overnight at 4C. Brains were carefully suctioned into a wide bore pipette tip and transferred to a cryomold. Excess 30% sucrose solution was removed via pipetting, and using a kim wipe. The brains were dried as much as possible, and manually manipulated using an eyelash into a region in the cryomold that was no bigger than the region that could be imaged when cryosectioned onto a slide. Individual brains were packed together, but not piled up, so that there was only a single layer of brains settled at the bottom of the cryomold. OCT was placed direction into the cryomold staring around the edges of the mold and moving inward to where the brains were placed to ensure even depositing. The brains rested in the OCT for 1 min at room temperature, ensuring no air bubbles were trapped between the tissue and the OCT, before being placed directly onto a slab of dry ice for rapid freezing. The cryomold was left on the dry ice for at least 10 mins before being stored in -80C until cryosectioning. The sample was attached to a chuck with OCT, and assembled in a cryostat set to -22C. The sample was carefully trimmed to a rectangular strip, and 8 micrometer cryosections were transferred to a coverslip. No trimming was used during cryosectioning. Test sections were checked before transfer to the coverslip and inspected under a microscope to confirm that sections through entire brains were obtained.

### Probe design

For the genome-wide labeling of metaloop anchors, we designed 73 10-kilobase probes centered on the anchor summits reported in Mohana Dorier Li et al. To trace the DIP-epsilon/DIP-zeta metadomain, we designed 92 5-kilobase probes tiling dm3 chr2L 6,300,000-6,450,000 and 8,950,000-9,150,000 and 9,400,000-9,550,000. To trace the sns/hbs metadomain, we designed X X-kilobase probes tiling dm3 chr2R4637505 - 4737505 and 10837505-11037505. Each probe is 100 bp long, with 20 bp complementary to fiducial probes, followed by 20 bp of barcode, then 40 bp of unique target sequence to bind the genome and 20 bp of a primer sequence used for probe synthesis. The probes were designed using software previously reported in Mateo et. al.^62^ and found at https://github.com/BoettigerLab/ORCA-public. The probe library was ordered from CustomArray (now operated by Genscript) as an oligo pool.

### Imaging procedure

Slides were first imaged in 2x SSC buffer to acquire GFP signal. DAPI was then added, and fields of view (FOVs) were acquired. Prior to ORCA hybridization, samples were removed from the imaging chamber. Following hybridization, samples were reassembled in the chamber, remounted, and the original GFP-imaged positions were relocated. DAPI was subsequently added again, and the same FOVs were reimaged.

### ORCA hybridization

We followed the procedure from Mateo et. al.^62^ (Hybridization with DNA primary probes, steps 147–166.

### Metaloop anchor ORCA image analysis

Nuclei in the *Drosophila* brain vary substantially in shape and size, and throughout much of the brain are densely packed together. This makes membrane based segmentation difficult and less reliable. Exploiting the specificity and uniqueness of our ORCA probes, and their distribution across all the large chromosomes, we used a DNA based segmentation approach for segmentation. In Drosophila, *homologous* chromosomes pair along their length during interphase, with the exception of a short interval, typically minutes to an hour, after cell division while pairing is re-established. Due to this pairing and the diffraction limit of fluorescent light, each metaloop anchor appears only once per nucleus (rather than as two separate homologs). DNA spots were assigned to cells using a seeded local 3D clustering approach. Spots detected in the first hybridization were used as provisional cell seeds, with seed pairs separated by less than 2 µm treated as unresolved and reduced to a single seed. For each seed and hybridization, up to ten nearest candidate spots were identified in 3D space. Candidate spots were retained only if they were within a 5-µm radius of the seed. These spots were used in the first iteration to provide a provisional list of cell positions. For each provisional cell, candidate spots from all hybridizations were pooled, and pairwise Euclidean distances between candidates were calculated in 3D. For each hybridization, the candidate that minimized the summed distance to the complete candidate set was selected as the representative spot for that cell. This procedure assigns at most one spot per hybridization to each cell while allowing hybridizations to be absent when no suitable candidate is detected. It ensures the assignments are maximally compact, as expected for genomic loci that share a convex boundary. The approach is conceptually similar to the minimization approach published by Le et al ^63^ to cluster unique DNA probe signals into separate chromosome territories for each homolog within a cell, and we built on that approach.

### Metadomain ORCA image analysis

Images were analyzed using the ChrTracer3 pipeline as described in Mateo et. al^62^.

### Metaloop anchor ORCA data analysis

#### Filtering

The data extracted from image processing was filtered in the following ways before proceeding with analysis. Cells with at least 95% of probes detected and with diameters of the spread of probes in x,y 6 micrometers or less were retained, while others were filtered out along with cells with no data. We then performed additional filtering to account for possible imaging artefacts that could arise from performing ORCA, such as false positive contacts due to failure of probe removal between rounds of imaging. We performed dimensionality reduction (PCA, then UMAP) on the probe-probe distance data and identified clusters of cells with close interactions between sequentially labeled probes. These filtering steps resulted in the 79,732 cells used in the analysis for the paper.

### Spatial analysis

Complete cross sections of the larval CNS were manually selected from the images for spatial analysis of metaloops at the tissue-scale. 5 regions were manually drawn to partition cross sections from the anterior to posterior side of the CNS (posterior towards the VNC), and from the periphery to the central brain. Regions with roughly equal width were drawn.

### Metadomain ORCA data analysis

#### Filtering

For the analysis in Figure 3, traces with at least 50% of the ORCA probes were selected. Traces that did not co-localize with the DAPI signal were removed. For the analysis in Figure 4-5, traces with at least 20% of the ORCA probes were selected, and traces that did not co-localize with the DAPI signal were removed as well.

#### Single-cell correlation with GFP

We computationally aligned images, using DAPI stains captured when the GFP imaging and ORCA was performed. This allowed us to then determine the GFP intensity at the 2D positions of each ORCA trace. A trace was assigned to be GFP+ if log10 of the GFP signal at that position was greater than 4, and GFP-if less than 1.

