## Supplementary Information for "Multiscale spatial analysis implicates chromosomal metaloops in gene patterning across the *Drosophila* brain"

| anchor ID | anchor chr | anchor summit | anchor type | nearest TSS | anchor distance to TSS (in bp) | Readout # |
|---|---|---|---|---|---|---|
| A1 | chr2L | 395,730 | intergenic | CG4213 | 6263 | 1 |
| A19 | chr2L | 17,260,843 | promoter | beat-IIIc | 60 | 2 |
| A20 | chr2L | 17,328,066 | intergenic | CG45691 | 7303 | 3 |
| A6 | chr2L | 2,595,538 | intergenic | CG15394 | 3363 | 4 |
| A7 | chr2L | 2,612,216 | intergenic | CG15395 | 2172 | 5 |
| A8 | chr2L | 2,677,718 | promoter | CG31690 | 0 | 6 |
| A4 | chr2L | 2,006,561 | promoter | CG33543 | 199 | 7 |
| A9 | chr2L | 4,793,528 | promoter | CG3294, fipi | 42, 0 | 8 |
| A5 | chr2L | 2,110,019 | promoter | dpr3 | 101 | 9 |
| A15 | chr2L | 10,964,468 | promoter | dpr2 | 45 | 10 |
| A10 | chr2L | 6,357,019 | intergenic | CG9500 | 1241 | 11 |
| A13 | chr2L | 9,464,720 | intergenic | numb | 14710 | 12 |
| A11 | chr2L | 6,411,319 | promoter | DIP-epsilon | 0 | 13 |
| A12 | chr2L | 9,083,116 | intergenic | Toll-4 | 854 | 14 |
| A14 | chr2L | 9,486,401 | intergenic | Gdi | 8291 | 15 |
| A16 | chr2L | 13,341,295 | intergenic | CG16826 | 6427 | 16 |
| A22 | chr2L | 19,606,108 | promoter | Lar, CG46244 | 115 | 17 |
| A25 | chr2L | 20,242,845 | intergenic | CG17571 | 14048 | 18 |
| A17 | chr2L | 13,349,544 | intergenic | CG16826 | 1715 | 19 |
| A21 | chr2L | 19,590,664 | intergenic | Lar | 3209 | 20 |
| A24 | chr2L | 20,230,040 | intergenic | CG17571 | 26842 | 21 |
| A18 | chr2L | 13,363,156 | intergenic | CG9377 | 1542 | 22 |
| A23 | chr2L | 20,121,154 | intergenic | CG10651 | 13631 | 23 |
| A26 | chr2L | 21,683,718 | promoter | nolo | 86 | 24 |
| A27 | chr2R | 8,686,664 | intergenic | Jon44E | 4671 | 25 |
| A33 | chr2R | 22,059,061 | intergenic | CG13500 | 1340 | 26 |
| A28 | chr2R | 8,715,879 | intergenic | PGRP-SC2 | 1024 | 27 |
| A32 | chr2R | 21,978,122 | intergenic | Gr58c | 1336 | 28 |
| A29 | chr2R | 8,797,965 | promoter | sns | 192 | 29 |
| A30 | chr2R | 15,011,084 | promoter | hbs | 35 | 30 |
| A31 | chr2R | 15,109,605 | intergenic | chn | 5579 | 31 |
| A34 | chr3L | 857,573 | promoter | dpr20 | 53 | 32 |
| A36 | chr3L | 1,997,964 | intergenic | CG42841 | 7646 | 33 |
| A37 | chr3L | 2,229,036 | intergenic | Oseg2 | 202 | 34 |
| A35 | chr3L | 866,595 | promoter | CG12502 | 55 | 35 |
| A38 | chr3L | 4,800,875 | intergenic | CG7509 | 4354 | 36 |
| A43 | chr3L | 7,013,951 | promoter | Mp | 100 | 37 |
| A39 | chr3L | 4,805,945 | intergenic | CG7509 | 593 | 38 |
| A42 | chr3L | 6,995,618 | intergenic | Mp | 3171 | 39 |
| A40 | chr3L | 6,623,897 | promoter | GluRIA | 81 | 40 |
| A45 | chr3L | 9,251,848 | promoter | GluRIB | 83 | 41 |
| A41 | chr3L | 6,653,057 | intergenic | eIF4E4 | 12518 | 42 |
| A46 | chr3L | 9,330,223 | intergenic | PGRP-LA | 4054 | 43 |
| A44 | chr3L | 7,643,734 | intergenic | CG7506 | 2035 | 44 |
| A50 | chr3L | 17,254,356 | intergenic | CG6497 | 367 | 45 |
| A51 | chr3L | 17,269,623 | intergenic | CG13723 | 5161 | 46 |
| A52 | chr3L | 18,743,288 | intergenic | CG6885 | 512 | 47 |
| A53 | chr3L | 20,004,878 | promoter | kug | 80 | 48 |
| A49 | chr3L | 12,868,180 | intergenic | CG10943 | 5193 | 49 |
| A54 | chr3R | 4,052,981 | intergenic | CG42402 | 24900 | 50 |
| A59 | chr3R | 7,780,279 | intergenic | CG18747 | 2743 | 51 |
| A58 | chr3R | 7,753,739 | intergenic | pyd3 | 3615 | 52 |
| A57 | chr3R | 5,150,695 | promoter | dpr16 | 149 | 53 |
| A61 | chr3R | 12,021,193 | intergenic | Spt3 | 3934 | 54 |
| A62 | chr3R | 12,094,732 | promoter | dpr17 | 77 | 55 |
| A60 | chr3R | 8,306,450 | intergenic | Sgt1 | 1005 | 56 |
| A71 | chr3R | 31,016,985 | promoter | 5-HT7 | 56 | 57 |
| A63 | chr3R | 23,539,077 | promoter | Ugt303B2 | 105 | 58 |
| A66 | chr3R | 28,672,385 | intergenic | Sid | 1636 | 59 |
| A64 | chr3R | 23,557,393 | promoter | beat-IV | 0 | 60 |
| A65 | chr3R | 28,654,770 | intergenic | CG31050 | 4250 | 61 |
| A67 | chr3R | 28,735,691 | intergenic | AstA-R2 | 9043 | 62 |
| A69 | chr3R | 29,344,728 | intergenic | CG45546 | 23272 | 63 |
| A70 | chr3R | 29,376,969 | promoter | Ptp99A | 197 | 64 |
| A68 | chr3R | 28,801,996 | intergenic | CheB98a | 4649 | 65 |
| A72 | chrX | 945,938 | promoter | CG3703 | 110 | 66 |
| A77 | chrX | 16,063,379 | promoter | dpr18, CG12395 | 54, 171 | 67 |
| A73 | chrX | 986,370 | intergenic | CG18823 | 3831 | 68 |
| A74 | chrX | 1,009,972 | intergenic | TfIIA-S-2 | 1517 | 69 |
| A75 | chrX | 5,433,300 | intergenic | CG15784 | 2532 | 70 |
| A78 | chrX | 17,611,656 | intergenic | X11L | 11619 | 71 |
| A76 | chrX | 13,358,960 | promoter | Syt12 | 0 | 72 |
| A79 | chrX | 19,348,857 | promoter | nAChRalpha7 | 0 | 73 |

**Supplementary Table 1: Details of regions labeled with ORCA**

ORCA readouts spanned 5kb upstream and downstream of the anchor summit. The anchor ID, chromosome arm the anchor is on, genomic position of the anchor summit, anchor type (intergenic or promoter), gene for the nearest TSS, and how far away that TSS is from the anchor summit are shown (anchor information from Supplementary Information in Mohana, Doreir, Li et. al.[1])

### Supplementary Fig. 1

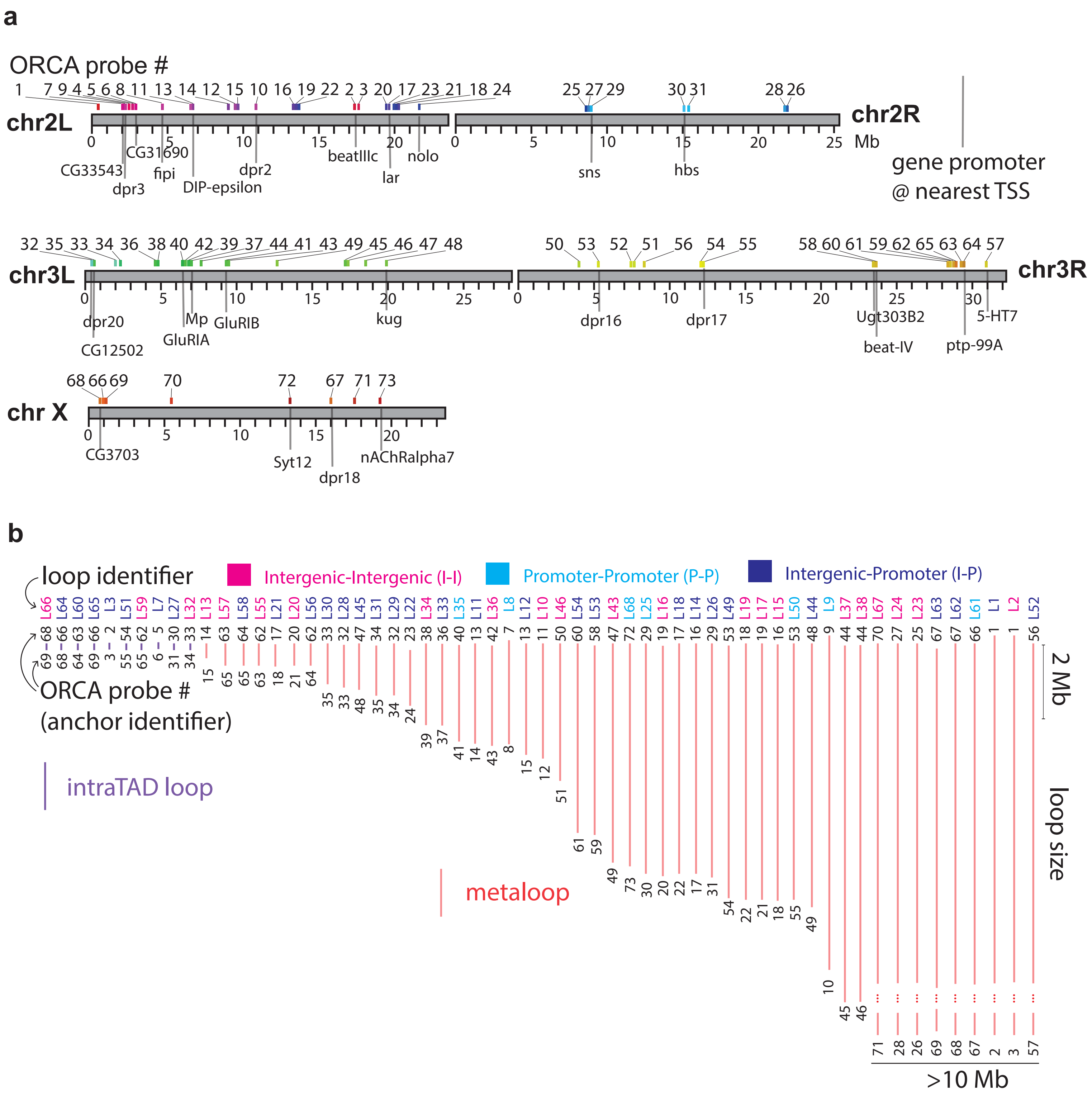

**Supplementary Fig. 1: Details of metaloop anchor labeling**

**a,** A schematic of the genomic positions of metaloop anchors that were labeled with ORCA probes (ORCA probes numbered 1-73). **b,** The loops formed by interactions between anchor pairs, categorized by intraTAD loops (shorter loop size) and metaloops (longer loop size), and whether the loops connected intragenic anchors to intragenic anchors (I-I), promoter anchors to promoter anchors (P-P), or intragenic anchors to promoter anchors (I-P).

#### Supplementary Fig. 2

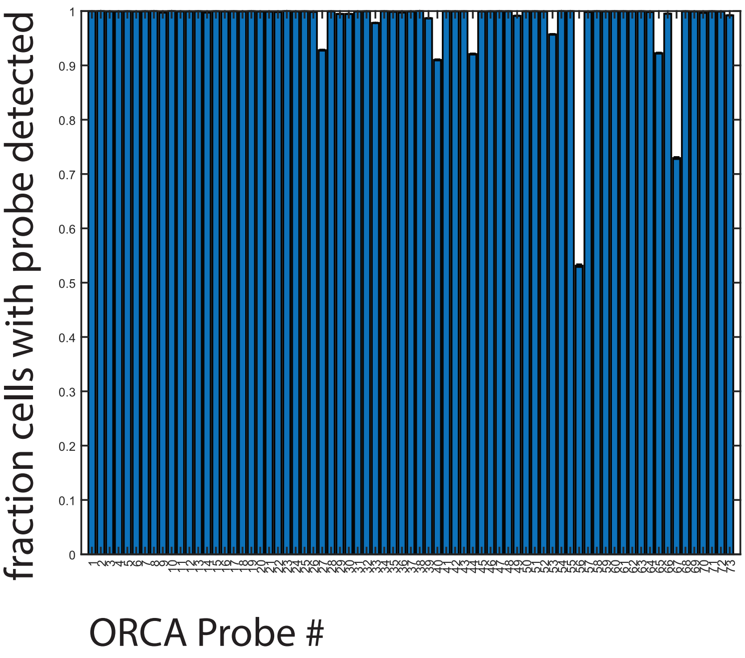

**Supplementary Fig. 2: Anchor detection efficiency**
Detection efficiency of each ORCA probe corresponding to a unique metaloop anchor.

### Supplementary Fig. 3

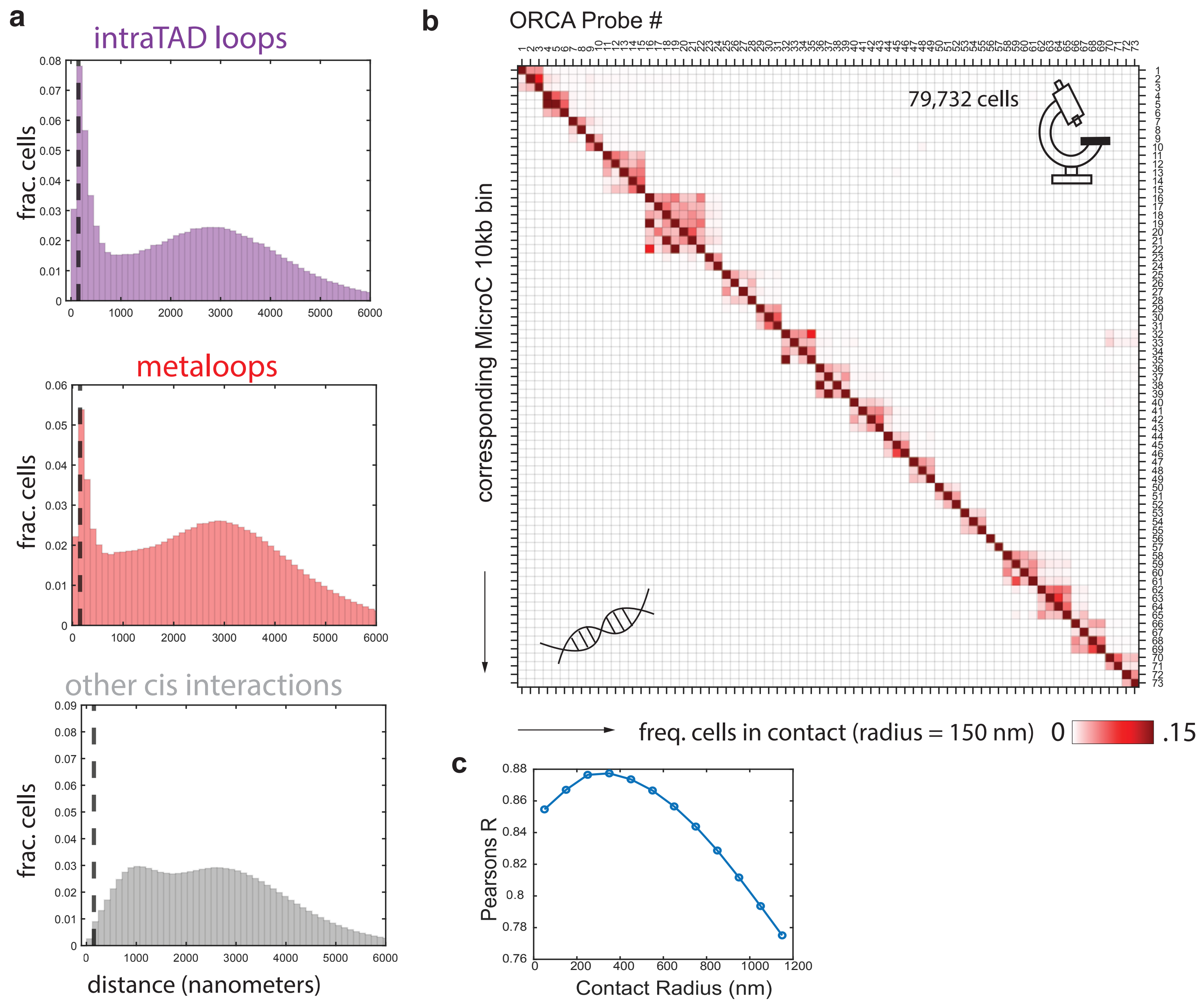

**Supplementary Fig. 3: Distance-based measurements of contact between metaloop anchors labeled with ORCA probes**

**a,** Distributions of the distances between pairs of probes corresponding to intraTAD loops, metaloops, and all other interactions occurring in *cis* on the same chromosome arm. The dotted line marks 150 nm. **b,** Contact maps generated from ORCA data (upper triangle) and Micro-C data (lower triangle) binned to 10 kb segments corresponding to the regions imaged with ORCA. Contacts were determined based on a 150 nm contact radius for the ORCA data. **c,** Correlation (Pearson's R) between contact maps generated from ORCA distance data thresholded based on various contact radii.

### Supplementary Fig. 4

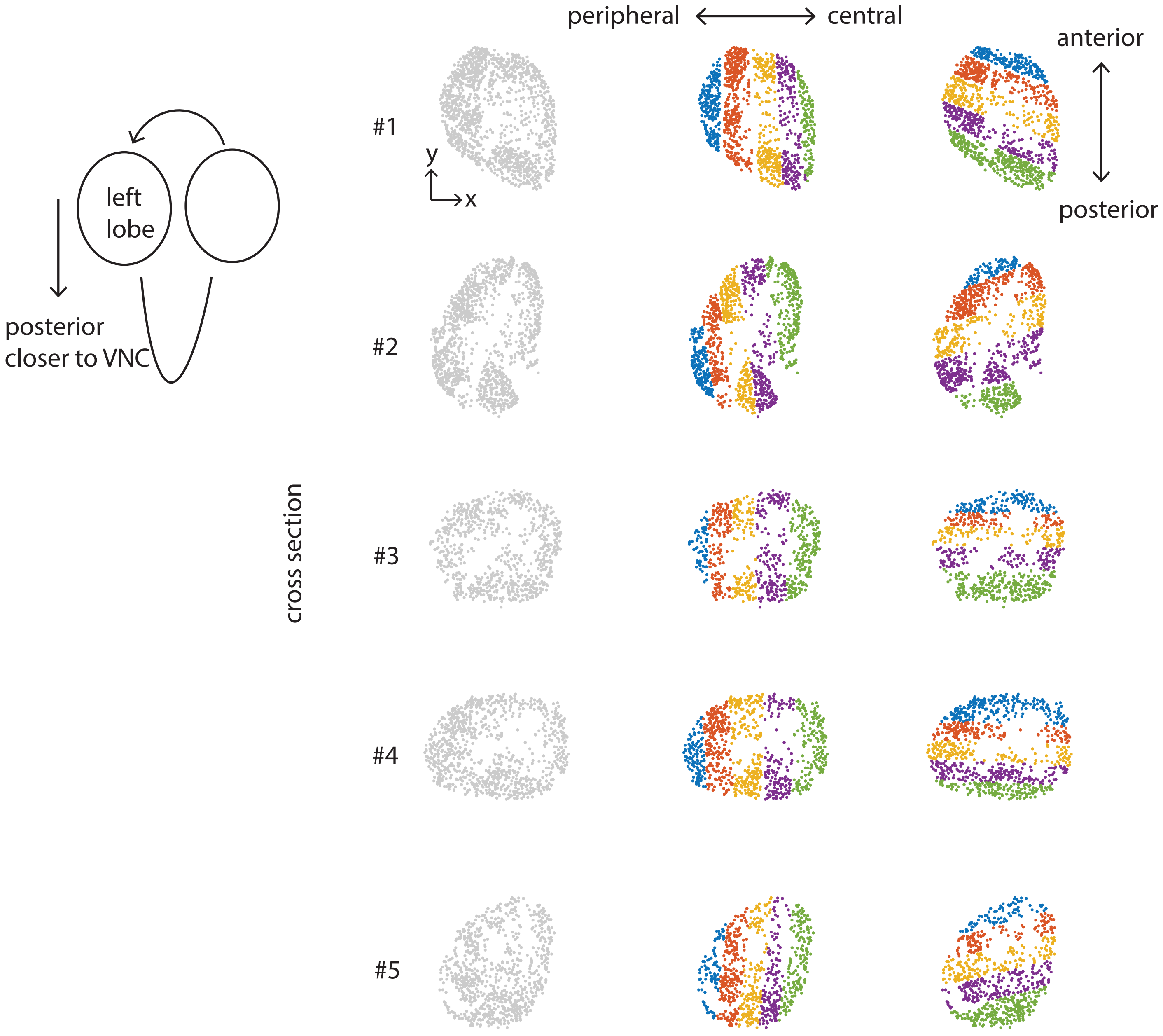

**Supplementary Fig. 4: Select cross section of the larval CNS segmented into pixels along the anterior-posterior and peripheral-central axes**

Cross sections were selected and oriented relative to the VNC. Cross sections were sub-divided from the peripheral to central and anterior to posterior parts of the brain. All cross sections were oriented with the periphery to the left of the central brain when the sub-regions were drawn.

#### Supplementary Fig. 5

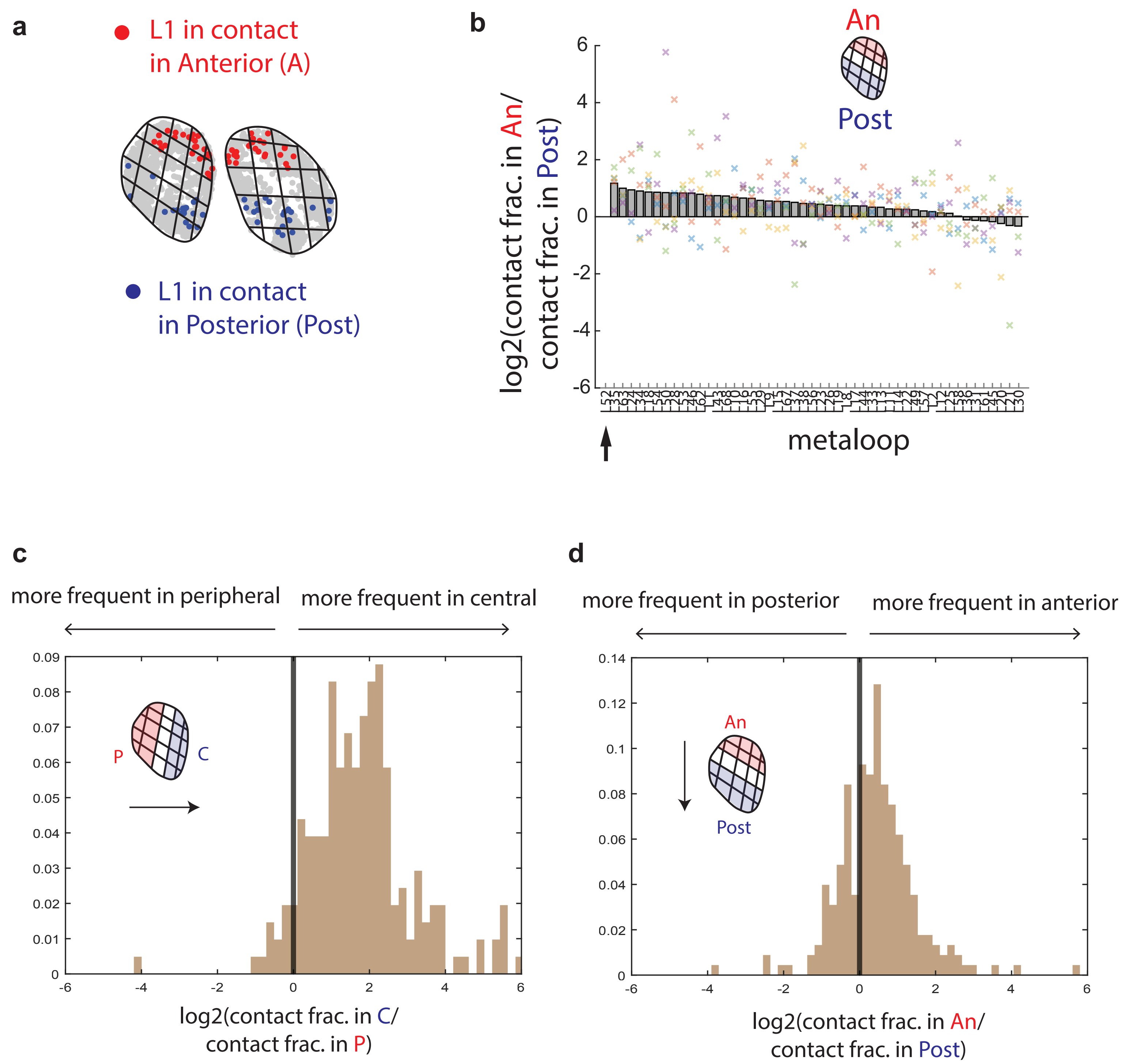

**Supplementary Fig. 5: Additional analyses of spatial biases of metaloop formation in the larval CNS**

**a,** Cells with L1 in contact in Anterior (An) pixels highlighted in red, and in Posterior (Post) pixels highlighted in blue. All other cells are grey. **b,** The log2 fold change in frequency of each metaloop in An versus Post, in 5 example cross sections, ranked by the mean. L52 is the top ranking metaloop (arrow). **c,** Histogram of the odds of any metaloop forming in the central versus peripheral region of the brain. **d,** Histogram of the odds of any metaloop forming in the anterior versus posterior region of the brain.

**Supplementary Fig. 6**

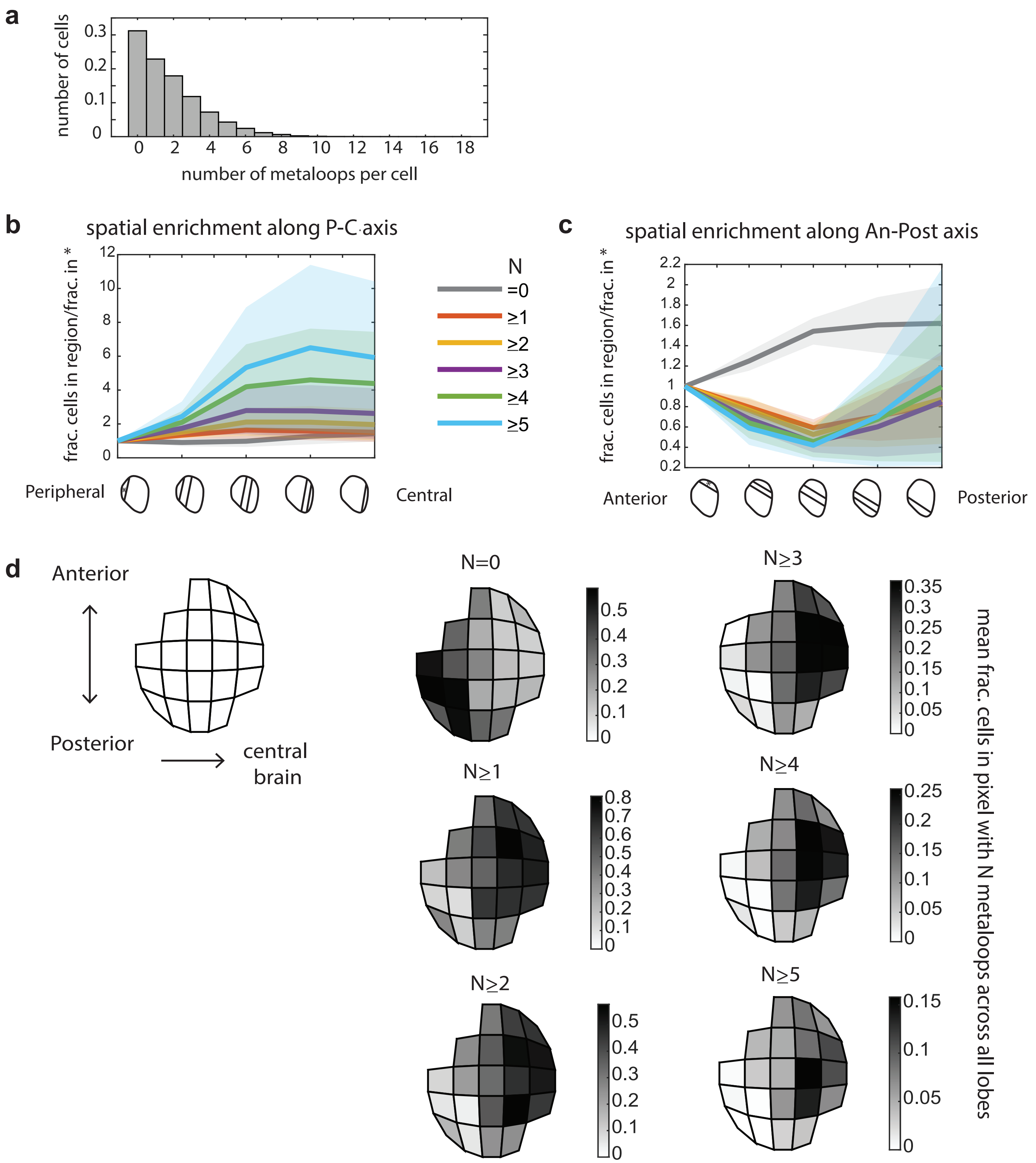

**Supplementary Fig. 6: Additional analyses of the spatial distributions of numbers of metaloops per cell across the larval CNS**

**a,** A histogram showing the distribution of the number of metaloops formed per cell. **b,** Enrichment of cells with more than N metaloops, calculated as the ratio of the fraction of cells in pixels aligned along the peripheral-central axis to the fraction in the most peripheral region. N = 1, 2, 3, 4 and 5. The line shows the mean in 5 cross sections, and the shaded region shows the standard deviation. **c,** The same as **b,** for pixels aligned along the Anterior-Posterior axis. **d,** Representations of the spatial patterns of how many metaloops are present per cell, averaged over 5 cross sections. Each pixel corresponds to an averaged spatial position across the independent cross sections. The mean fraction of cells in a spatial pixel with N metaloops is shown.

### Supplementary Fig. 7

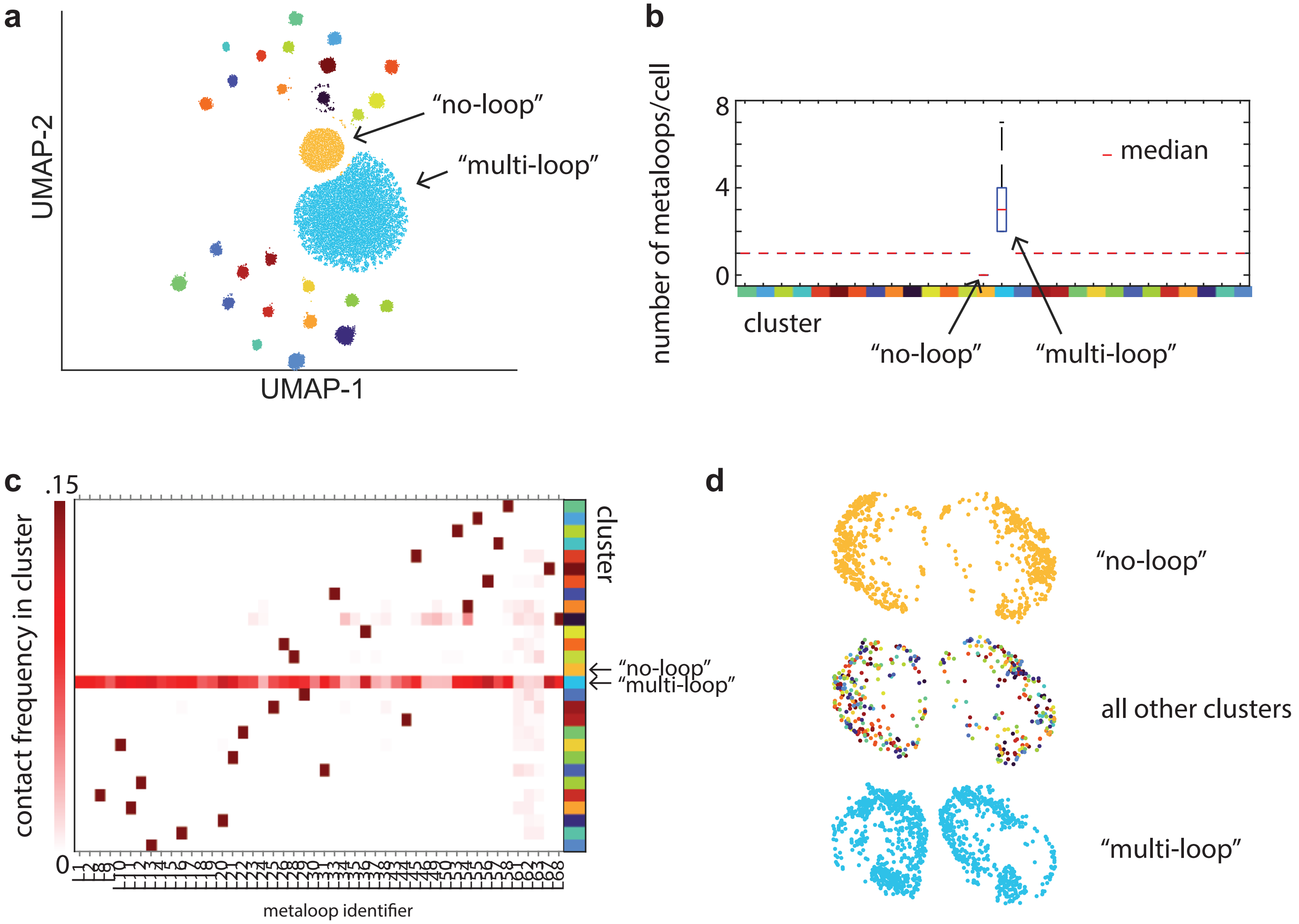

**Supplementary Fig. 7: Cells cluster together based on metaloop composition**

**a,** UMAP shows clusters of cells grouped by metaloop composition after performing dimensionality reduction on single-cell contact data. One cluster contains cells with no metaloops (“no-loop”) and another is named the “multi-loop” cluster based on the median number of metaloops per cell shown in **b**. **c,** Contact frequencies of metaloops within each cluster of cells. **d,** The xy positions of cells in the “no-loops” “multi-loop” and all other clusters are shown in an example cross section.

### Supplementary Fig. 8

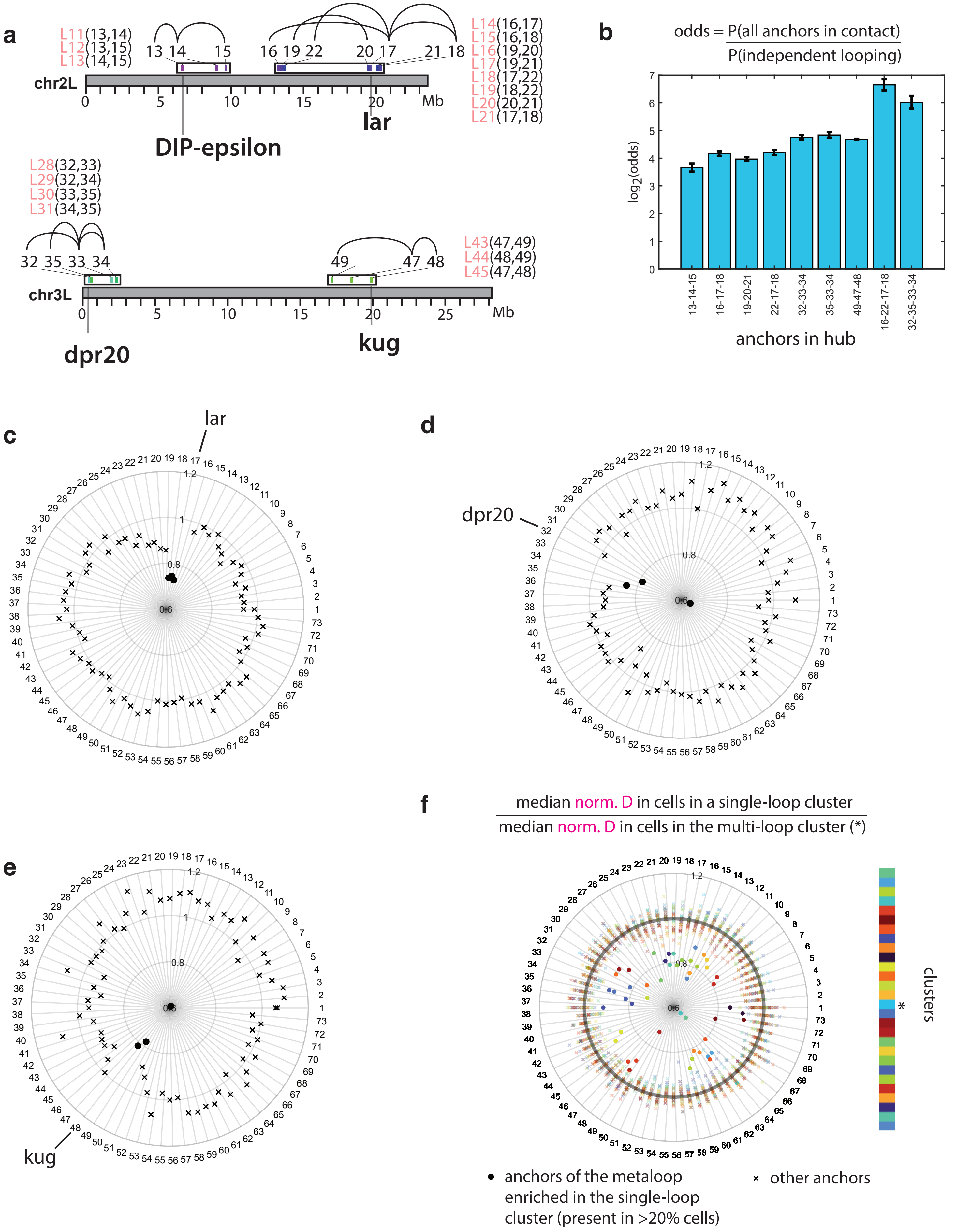

**Supplementary Fig. 8: Odds of formation and nuclear positioning of hubs**

**a,** Schematic of the anchors and metaloops that form hubs around a promoter anchor (indicated by the genes: *dpr20*, *kug*, *DIP-epsilon*, and *lar*). Arcs indicate which anchors are connected by metaloops. The metaloop identifier and corresponding anchors are labeled. **b,** The odds of multi-way contact for various hubs involving many anchors, indicated by the ORCA probe. Plots in **c,d,e,** show the median normalized distances to the geometric center of all anchors for cells with versus without the *lar*, *dpr20*, and *kug* hubs. Distances are normalized by the radius of gyration (Rg) of anchors in the cell. The x's show the ratio of normalized distances in hub vs. no hub cells for anchors that don't participate in the hub, while the filled dots show the ratios for anchors in the hub. **f,** A similar plot showing the ratios of normalized distances to the geometric center of all anchors calculated in individual clusters identified during dimensionality reduction. The ratio compares normalized distances found in the clusters, versus the multi-loop cluster (*), in which no single metaloop is particularly frequent. The filled dots show the ratios of normalized distances for anchors that are frequently in contact (>20% cells in the cluster), and x's show the ratios of normalized distances for all other anchors.

### Supplementary Fig. 9

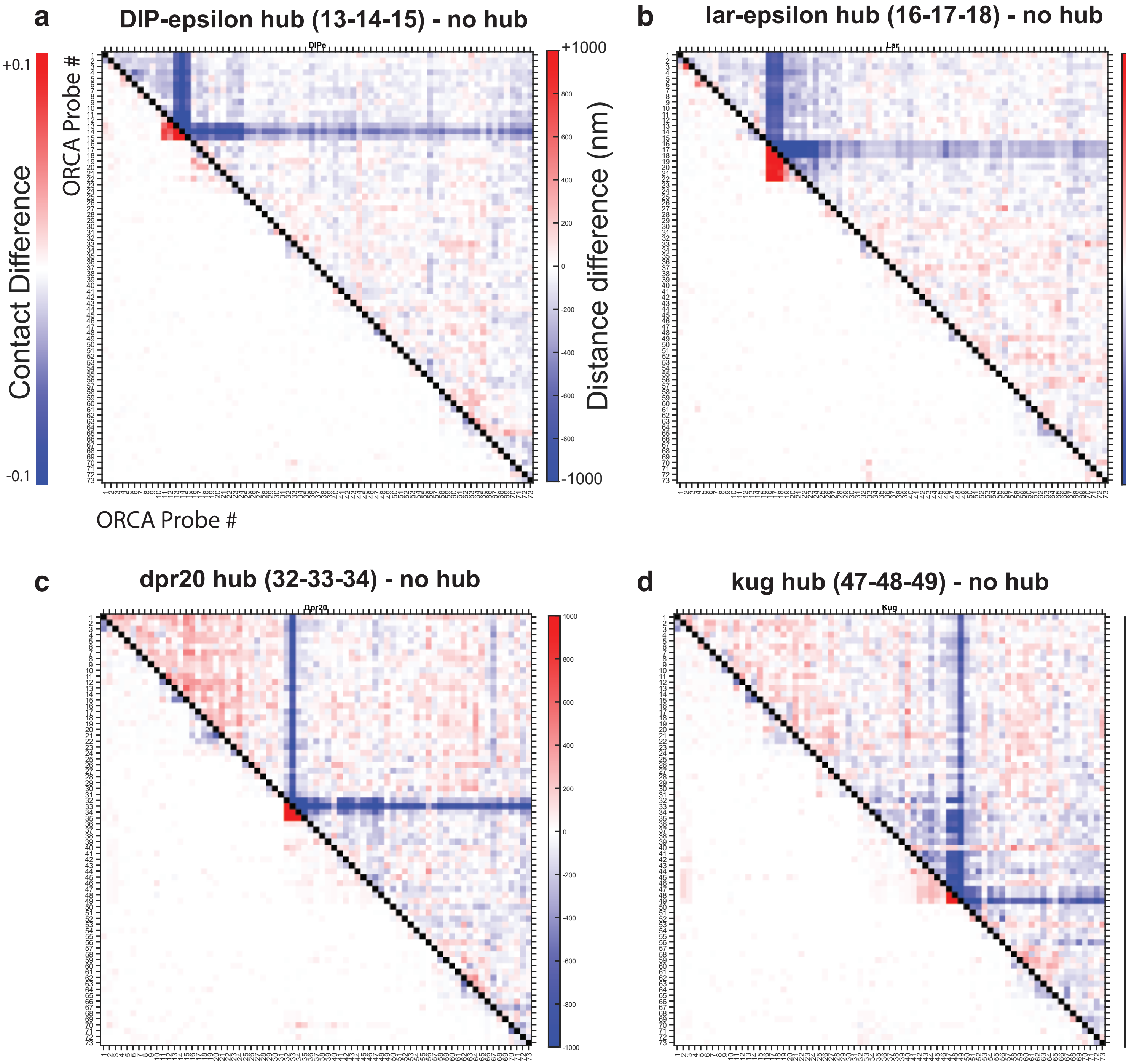

**Supplementary Fig. 9: Differences in distance and contact in cells with and without hubs**
**a,** The difference between the contact maps generated from cells with and without the *DIP-epsilon* hub are shown on the bottom triangle, and the difference between the distance maps is shown in the top triangle. The same is shown for the difference between cells with and without the *lar*, *dpr20*, and *kug* hubs are shown in **b,c,** and **d** respectively.

### Supplementary Fig. 10

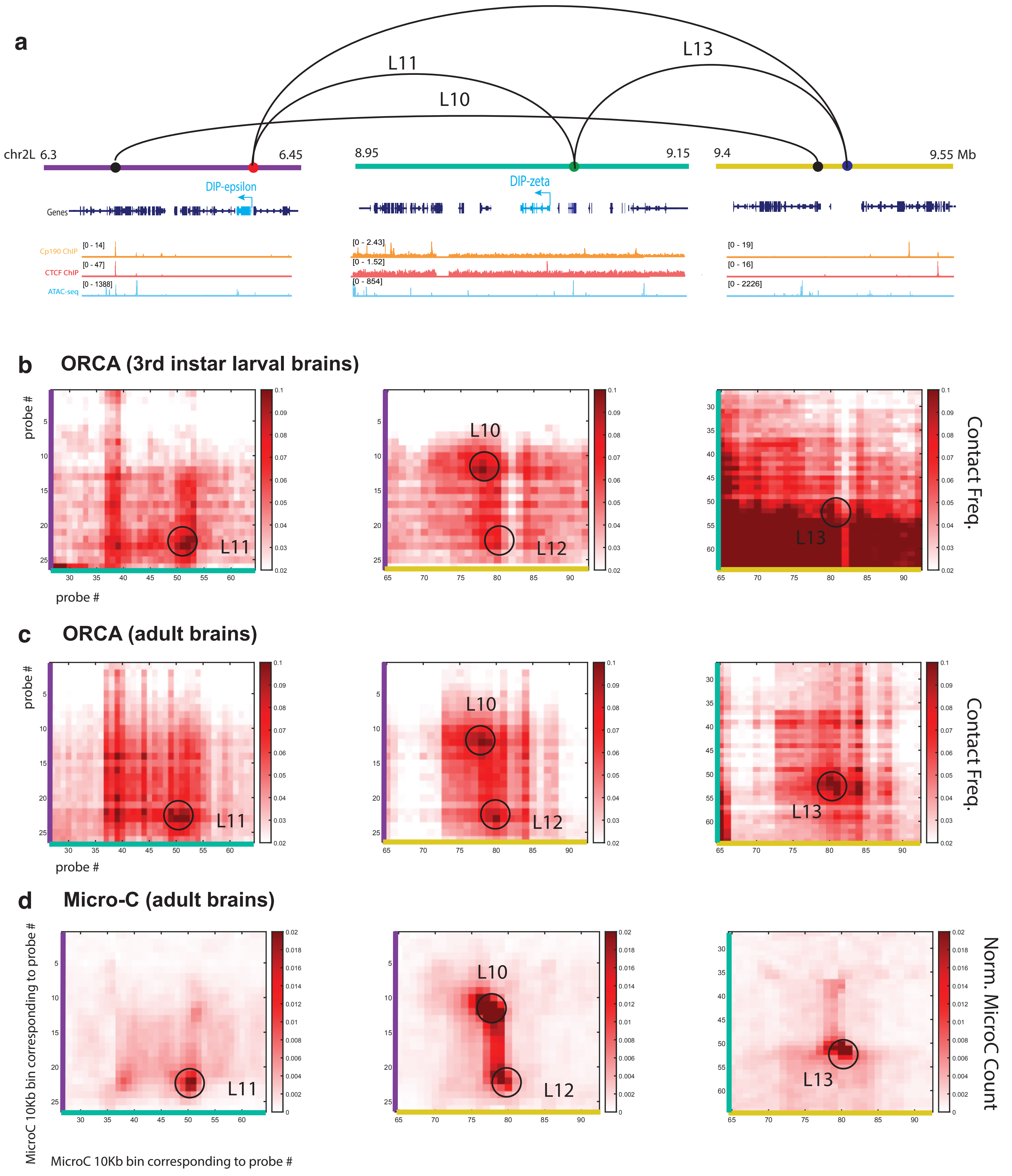

**Supplementary Fig. 10: Contact maps of metadomains captured in larval and adult brains**
**a,** Gene, ChIP-seq[2] and ATAC-seq[3] tracks below the regions traced with ORCA probes. Metaloop anchors connected by metaloops included in this tracing are shown. **b,** Contact maps showing the interactions between 3 distal regions spanning the anchors for metaloops L10,L11,L12, and L13. ORCA tracing was performed in the 3rd instar larval brain. **c,** The same contact maps for tracing performed in the adult brain are shown. **d,** Micro-C maps corresponding to the regions shown in the ORCA contact maps, binned to 5 kb, the size of the ORCA probes.

### Supplementary Fig. 11

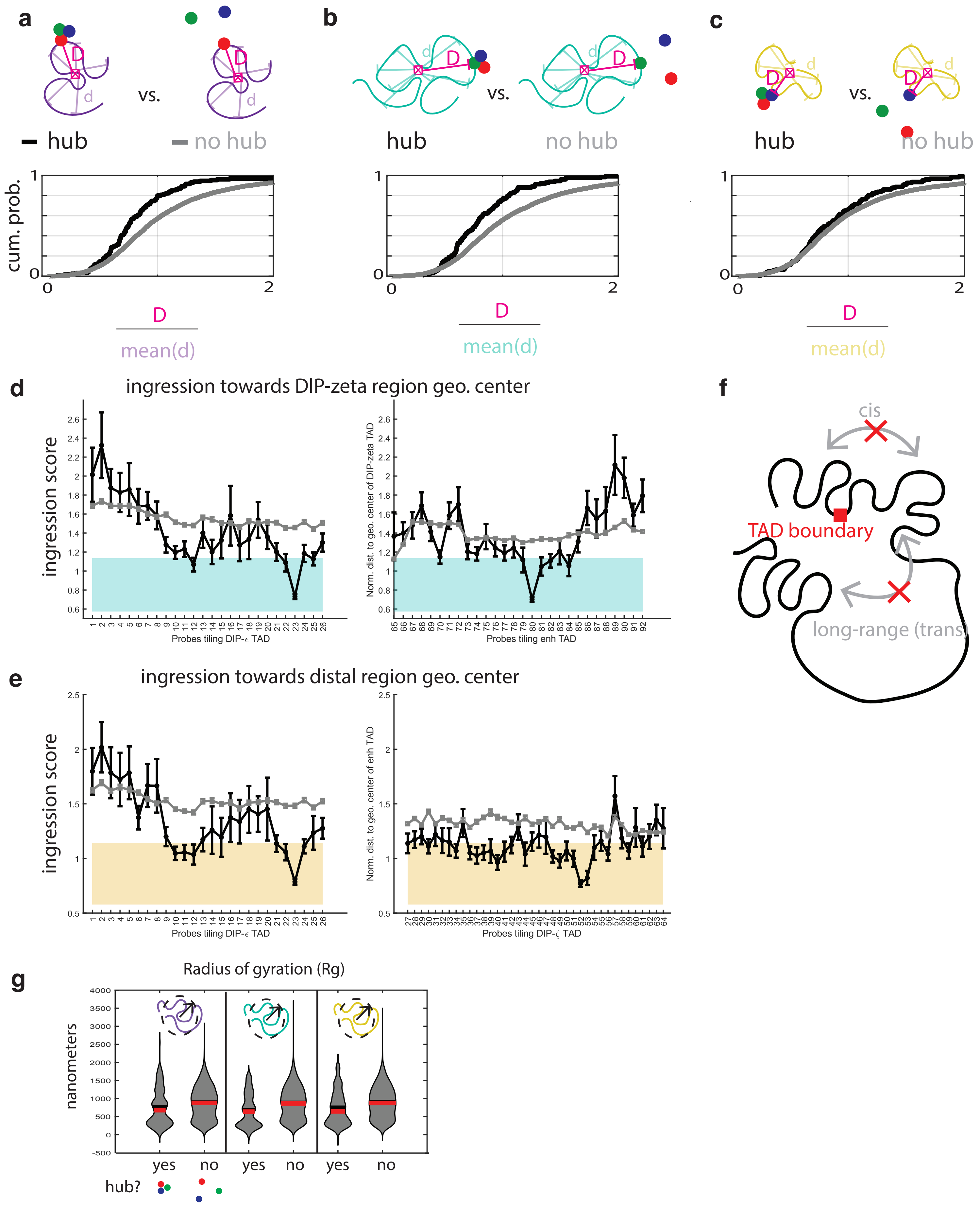

**Supplementary Fig. 11: Additional analyses of centering within chromosome segments traced with ORCA probes**

**a,** Cumulative probability distribution plots of the distances between the *DIP-epsilon* proximal anchor and the geometric center of the probes tiling the *DIP-epsilon* region in cells with and without the *DIP-epsilon* hub. Distances are normalized by the mean of the distances between all probes tiling the *DIP-epsilon* segment in that cell. **b,** Similar cumulative probability distribution plots for the distances between the *DIP-zeta* proximal anchor and the geometric center of the probes tiling the *DIP-zeta* region. **c,** Similar cumulative probability distribution plots for the distances between the anchor in the enhancer region and the geometric center of the probes tiling the enhancer region. **d,** As in Fig. 3, distances to the geometric center of the *DIP-zeta* region were measured on a probe-by-probe basis, normalized, and reported as ingression scores. The mean and s.e.m. of these ingression scores in traces with (grey) and without the *DIP-epsilon* hub (black). The shaded region marks the 25-75% confidence interval for all pooled distances between probes tiling the *DIP-zeta* segment and their geometric center. **e,** Similar measurements of the distances to the geometric center of the distal region. The shaded region marks the 25-75% confidence interval for all pooled distances between probes tiling the distal region and their geometric center. **f,** A schematic of the cis and long-range (trans) interactions mediated by boundary elements. **g,** Radius of gyration of the probes tracing each segment, in populations of cells with and without the hub. Distribution plots are shown.

### Supplementary Fig. 12

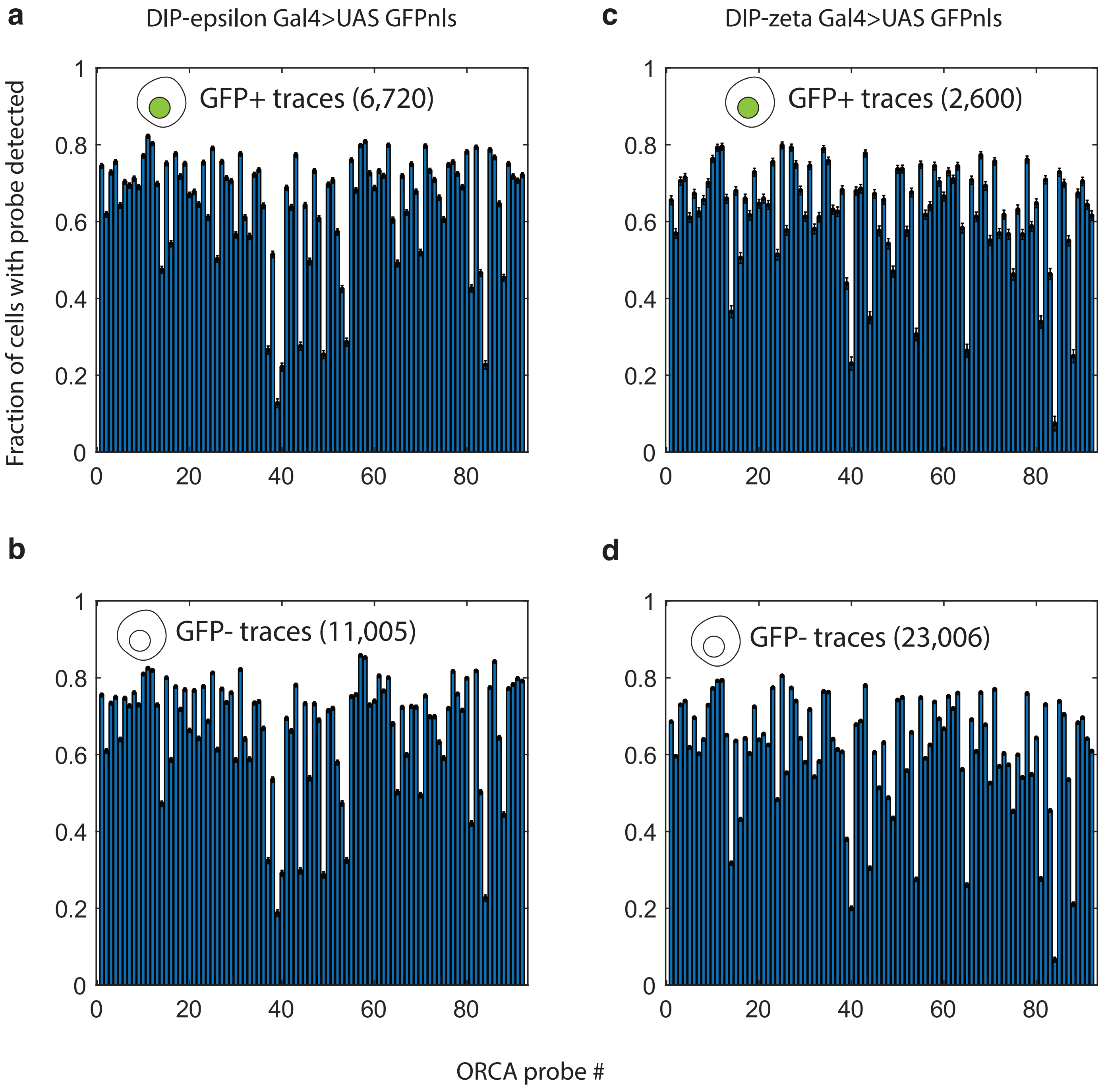

**Supplementary Fig. 12: Detection efficiencies in cells assigned an expression state based on GFP intensities**

**a,** Detection efficiency of ORCA probes in DIP-epsilon expressing cells. **b**, Detection efficiency in DIP-epsilon silent cells. **c,** Detection efficiency of ORCA probes in DIP-zeta expressing cells. **d,** Detection efficiency of ORCA probes in DIP-zeta silent cells.

### Supplementary Fig. 13

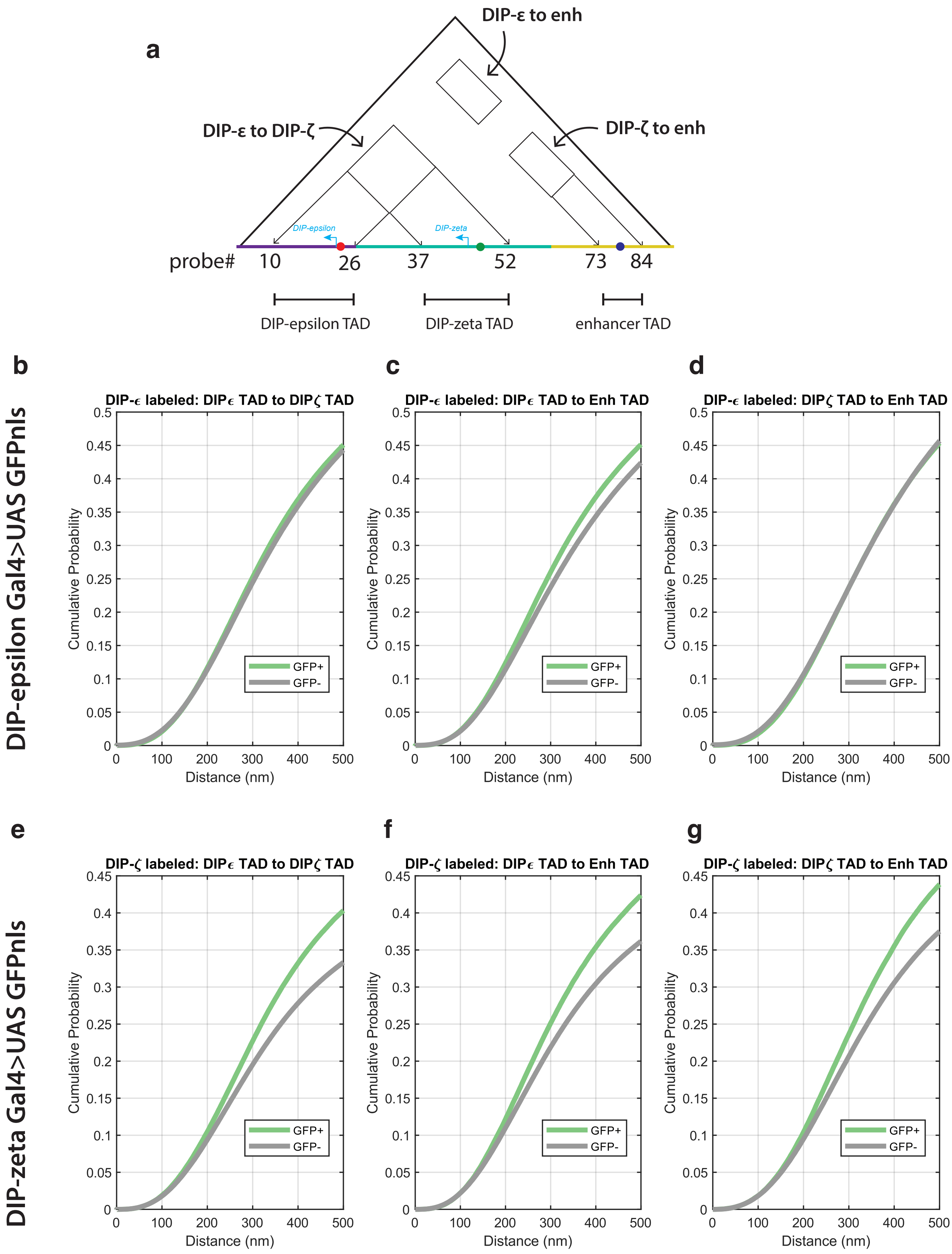

**Supplementary Fig. 13: Distributions of expression-dependent distances between distal TADs**

**a,** A schematic showing which probes span the TADs that interact in metadomains. TADs are named “*DIP-epsilon*”, “*DIP-zeta*” and “enhancer”. **b,** Cumulative probability distribution plots of all 3D distances (nanometers) from probes in the DIP-epsilon TAD to the DIP-zeta TAD in cells expressing GFP (marking *DIP-epsilon*) and not expressing GFP. **c,** The same for 3D distances from probes in the *DIP-epsilon* TAD to the enhancer TAD. **d,** The same for 3D distances from probes in the *DIP-zeta* TAD to the enhancer TAD. **e,f,g,** show similar measurements, except GFP instead labels *DIP-zeta* expression rather than *DIP-epsilon*.

### Supplementary Fig. 14

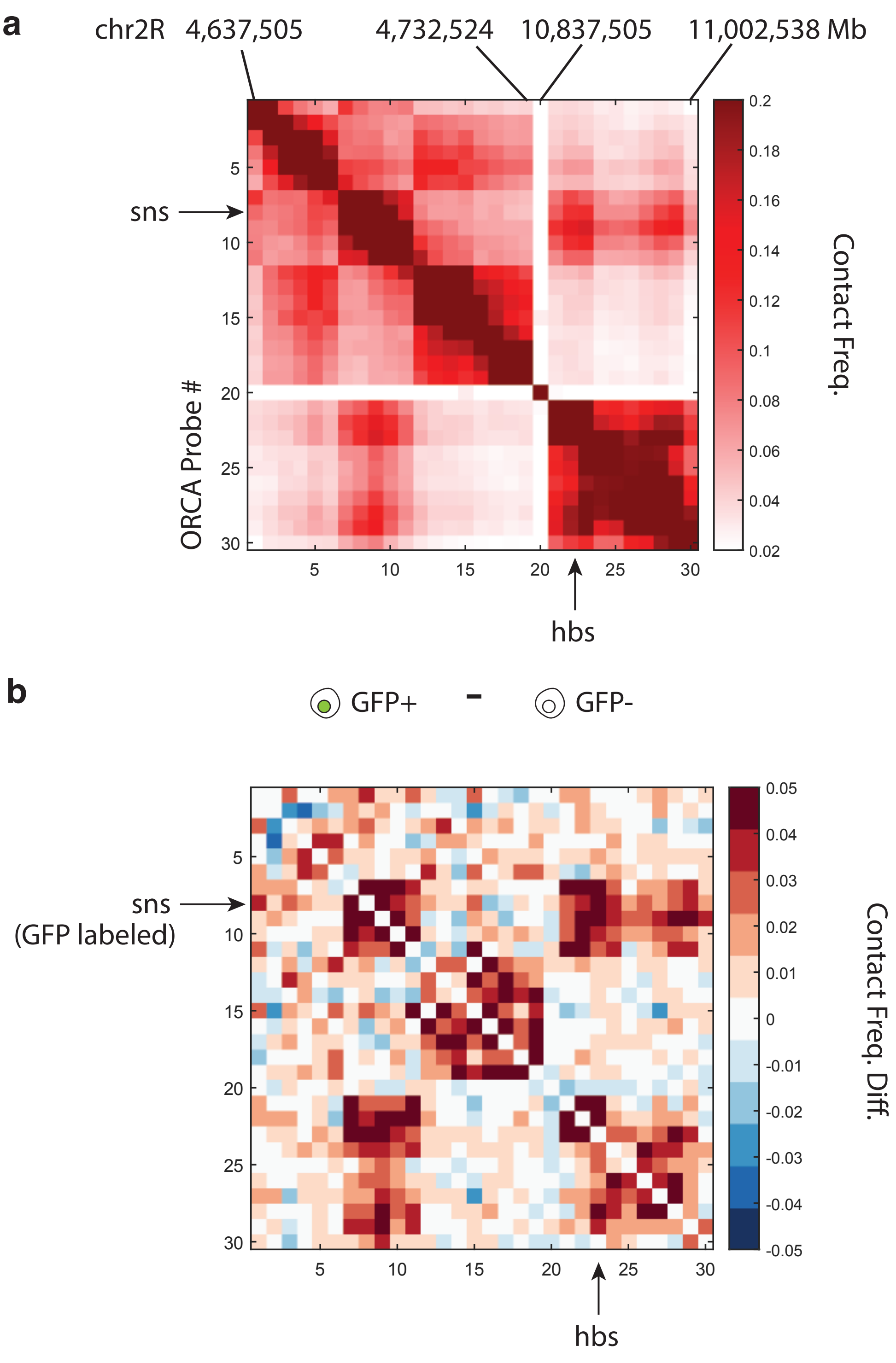

**Supplementary Fig. 14: Increased contacts between TADs in *sns*-expressing cells in the larval CNS**

**a,** A contact map of the metadomain spanning the genes *sns* and *hbs.* **b,** The differences in pairwise contact frequencies between *sns*-expressing GFP+ cells and GFP- cells.

### Supplementary Fig. 15

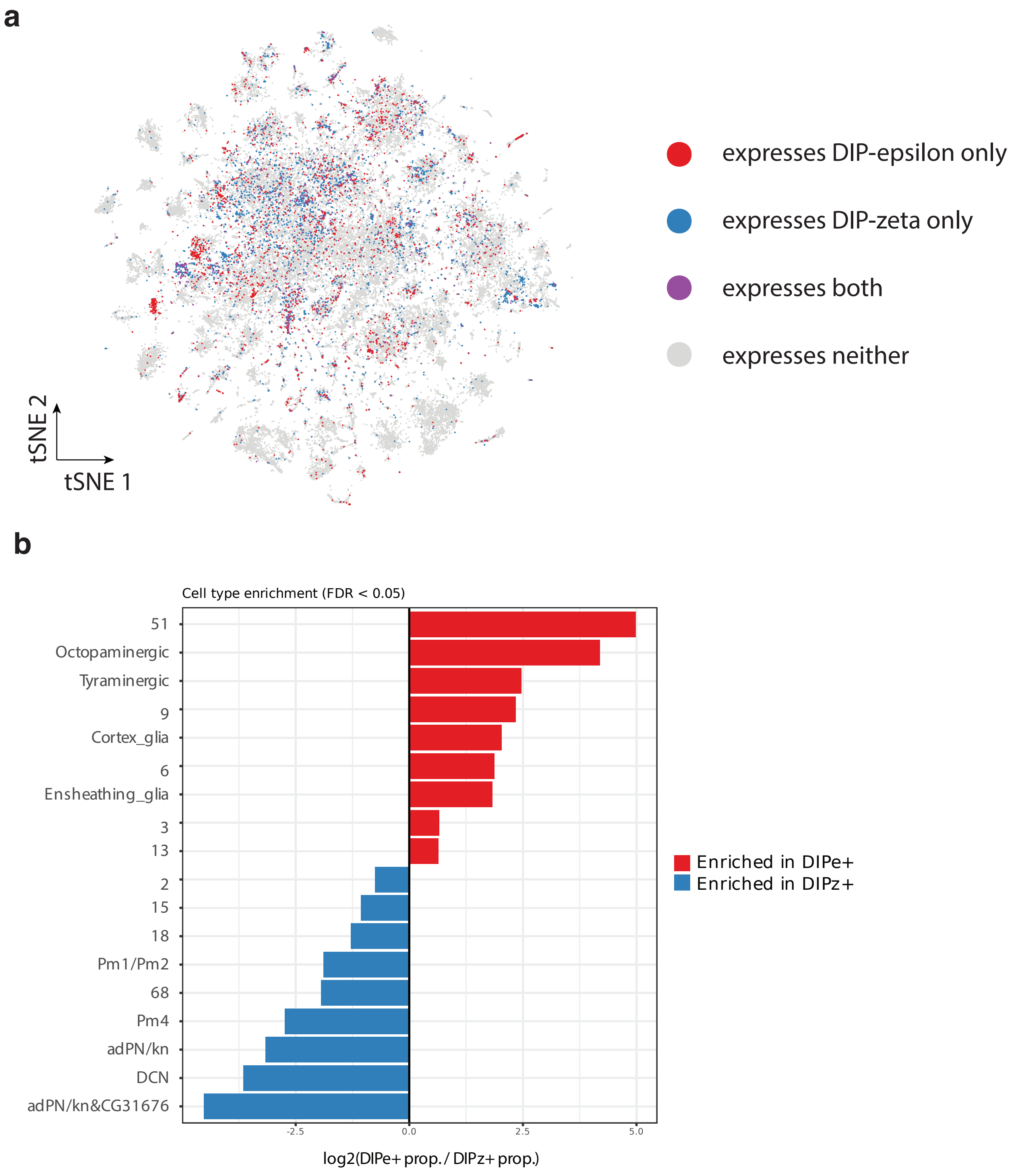

**Supplementary Fig. 15: *DIP-epsilon* and *DIP-zeta* expression in single cells**

**a,** tSNE embedding of cell types mapped by published scRNAseq in the Drosophila adult brain[4].

**b,** Enrichment of *DIP-epsilon* and *DIP-zeta* expression in different adult brain cell types.

### Supplementary Fig. 16

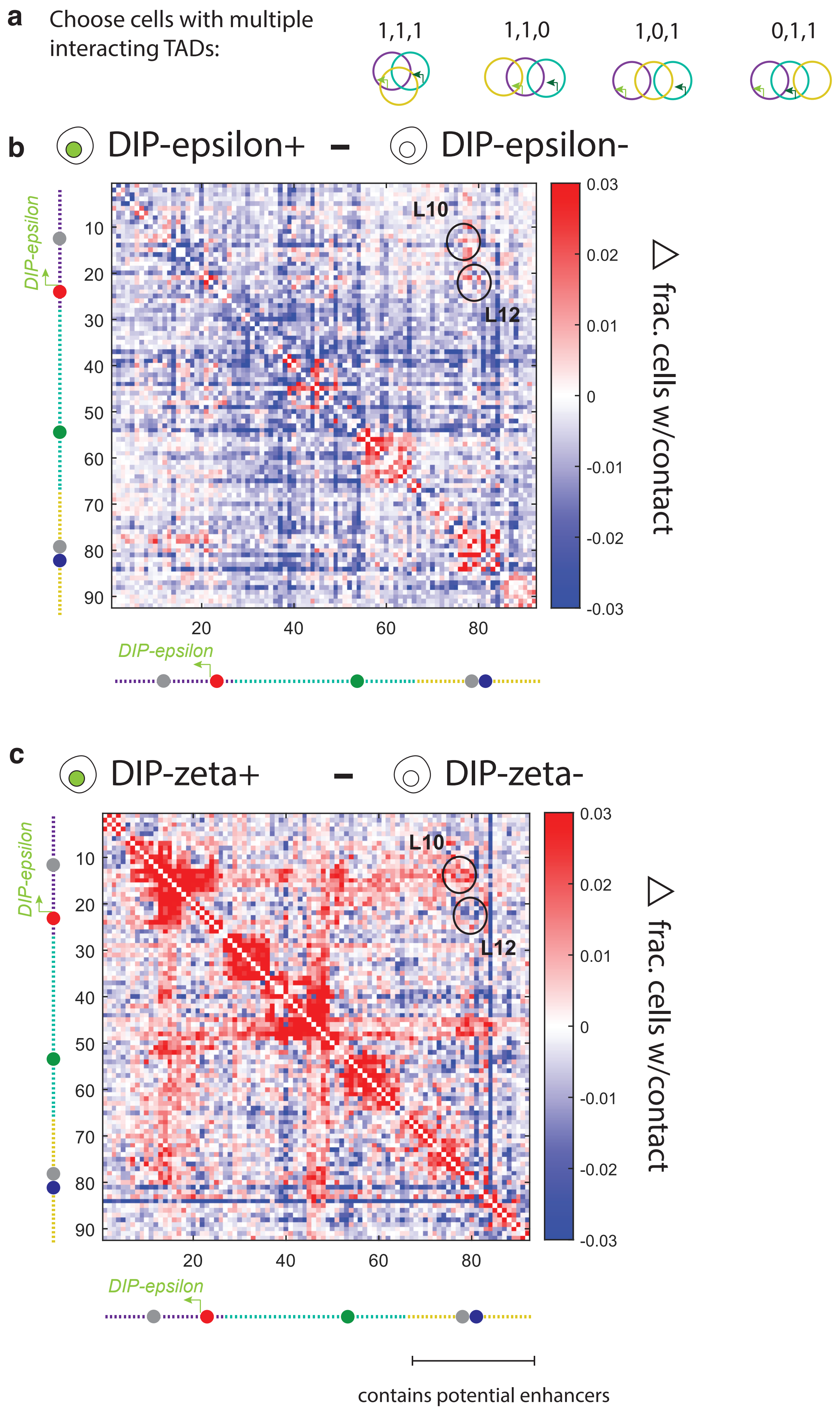

**Supplementary Fig. 16: Expression dependent contact differences in sub-populations of cells harboring multi-TAD interactions**

**a,** We selected cells with multiple interacting TADs for further contact analysis. **b,** The differences in pairwise contact frequencies between cells that are expressing *DIP-epsilon* (GFP+) and GFP**-** in after selection. **c,** The contact differences between cells that are expressing and not expressing *DIP-zeta*.
